# Investigation of plant virus-like particle formation in bacterial and yeast expression systems

**DOI:** 10.1101/2024.09.20.614045

**Authors:** Ina Balke, Gunta Resevica, Vilija Zeltina, Ivars Silamikelis, Elva Liepa, Reinis Liepa, Ieva Kalnciema, Ilze Radovica-Spalvina, Dita Gudra, Janis Pjalkovskis, Janis Freivalds, Andris Kazaks, Andris Zeltins

**Author notes:** Corresponding authors (I. Balke); (A. Zeltins).

## Abstract

Virus-like particles (VLPs) have garnered significant attention due to their potential applications in various fields, particularly in biomedicine, where they are used for drug delivery, vaccine development, and as diagnostic tools. Plant virus-derived VLPs have been especially successful, leading to a surge in research aimed at developing new VLPs. Despite this growing interest, not all attempts at VLP formation are successful. Understanding the factors that contribute to successful VLP assembly is crucial for advancing this field. In this study, we focus on the coat proteins (CPs) of three plant viruses belonging to the Sobemovirus genus as a model to investigate the process of VLP formation. Our findings demonstrated that the strong binding of CPs to ssDNA can be a major reason for unsuccessful VLP production. This issue can potentially be overcome by transitioning from an episomal to a chromosome-integrated expression system. By doing so, we aim to elucidate the mechanisms that facilitate successful VLP assembly. Our findings provide valuable insights into the properties of sobemovirus CPs and highlight their potential for encapsulating foreign nucleic acids, thereby providing additional applications as nanocontainers or vaccine platforms with interior modification capabilities for enhanced immune response. Moreover, our RNA-Seq data indicates that sobemovirus-derived VLPs predominantly package their CP mRNA, irrespective of the expression system used. This characteristic can be utilized for the functional encapsulation of nucleic acids, enhancing the versatility and utility of these VLPs in various biotechnological applications.

## 1. Introduction

The development of gene engineering techniques has significantly advanced the fields of biology, biotechnology, medicine, and vaccinology. These advancements have led to the creation of virus-like particles (VLPs) from cloned viral structural genes expressed in heterologous host cells, culminating in the first recombinant subunit vaccine (for review, see [1]). This breakthrough paved the way for a new class of subunit vaccines, resulting in the approval of five VLP-based vaccines for human use: hepatitis B [2], cervical cancer (Gardasil and Cervarix [3-5]), hepatitis E (Hecolin [6, 7]), malaria (Mosquirix [2, 8, 9], and COVID-19 (Covifenz [10]). Extensive research on plant viruses over the past century has provided profound insights into viral capsid structures, their assembly mechanisms, and lifecycles. This knowledge has been applied biotechnologically to develop model vaccines and versatile nanoparticle platforms based on more than 55 different plant viruses [1, 11, 12]. These studies have shown that plant virus-derived VLP platforms can serve as multipurpose tools. Plant virus-derived nanoparticles have several advantages over those derived from mammalian viruses. They typically demonstrate little to no pre-existing immunity and offer structural flexibility for various manipulations due to their composition, usually involving only one or a few coat protein (CP) molecules [11]. All VLPs lack replicative viral nucleic acid (NA) but mimic the structure of the original virus. Depending on the expression system, they can encapsulate toll-like receptor (TLR) ligands such as TLR7/8, TLR9, TLR3, or non-TLR binding ligands to enhance immune responses [13-17]. The geometric structure and repetitive surface of VLPs mimic pathogen-associated molecular patterns (PAMPs) and pathogen-associated structural patterns (PASPs), which can promote robust antibody responses [18-20]. VLPs can display high-density target antigens through various chemical and genetic fusion techniques. Their typical size of 20-200 nm facilitates efficient lymph node drainage and interaction with antigen-presenting cells (APCs) and B cells [21, 22]. Plant virus CPs can be expressed in various systems [1, 12, 23]. They are stable and suitable for large-scale production [24, 25]. These features make plant virus-derived VLPs promising candidates for diverse biomedical applications, including vaccine development and nanoparticle platforms. Sobemoviruses are small, positive-sense RNA viruses that encode a single coat protein (CP) molecule translated from subgenomic RNA. The virion diameter ranges from 26 to 32 nm, consisting of 180 CP monomers arranged in a *T = 3* lattice symmetry [26]. The 3D structures of five sobemoviruses have been elucidated: southern cowpea mosaic virus (SCPMV, previously known as the southern bean mosaic virus C-strain) at 2.8 Å resolution [27], southern bean mosaic virus (SBMV, known as the SBMV B-strain) at 2.9 Å resolution [28], Sesbania mosaic virus (SeMV) at 4.7 Å [29] and 3 Å resolutions [30, 31], rice yellow mottle virus (RYMV) at 2.8 Å resolution [32], cocksfoot mottle virus (CfMV) at 2.7 Å [33], and ryegrass mottle virus (RGMoV) at 2.9 Å resolution [34]. These structures reveal a canonical jellyroll beta-sandwich fold and are nearly identical. Notable differences were observed in RGMoV, where the absence of the FG loop resulted in the loss of interaction between the loop and the beta-annulus, compensated by additional interactions between the N-terminal arms, reducing the RGMoV diameter by 8 Å [34]. Self-assembled VLPs of SeMV, CfMV, and RYMV CPs have been successfully produced using bacterial expression systems [23, 35, 36]. SeMV CP forming VLPs have been used in structural studies [37-42], Additionally, sobemoviral VLPs have been formed in different lattice symmetries – *T = 1* [43-46] and *T = 2* [35], and β-annulus motif assembly in nanocages [47, 48]. SeMV nanoparticles has been used as *in vivo* imaging agents [49], examined for the biodistribution, toxicity and histopathological changes [50], and tested for genetic incorporation of antigen sequences and used for intracellular delivery of antibodies [51]. CfMV CP has been tested for foreign epitope insertion [36]. These experiments collectively highlight the potential of sobemoviruses as a safe and flexible platform for various applications, including vaccine development, imaging, and delivery agents.

The bacterial expression systems based on *Escherichia coli* are well-characterized, with various commercial expression vectors available, making it the most popular platform for recombinant protein production. Compared to yeast, mammalian, and insect cell cultures, *E. coli* offers several advantages, including rapid growth kinetics, the ability to achieve high cell density cultures, and the use of readily available and inexpensive media components with rich complex additives. Additionally, transformation is fast and easy in *E. coli* [52]. The demand for low-cost production of complex recombinant proteins and therapeutics for clinical applications has driven the development of specialized *E. coli* strains. For example, the endotoxin-free *E. coli* strain ClearColi^™^ BL21(DE3) was developed to enable protein expression without endotoxin contaminations [53]. The SHuffle^®^ T7 strain is designed for the correct folding of disulfide-bonded proteins [46], and a system for producing glycosylated proteins was developed based on this strain [54, 55]. Other examples include the LOBSTR strain for low-expressing protein purification with reduced background contamination [56], and the Lemo21(DE3) strain for membrane protein overexpression [57]. Despite the availability of multiple *E. coli* expression systems, not all expressed plant virus CPs form VLPs. Some CPs, such as those from cowpea chlorotic mottle virus (CCMV) and cucumber mosaic virus (CMV), can be assembled *in vitro* from purified inclusion bodies [58, 59] or *in E.coli* cells during the cultivation by simple reducing the temperature during CP expression [22, 36]. Some reports suggest a change to a yeast expression system if the VLP formation in *E. coli* cells is not achieved [60, 61].

Despite various manipulations, the *E. coli* expression systems are not generally recognized as safe (GRAS), unlike yeast systems [62]. The *Pichia pastoris* system (recently reclassified as *Komagataella phaffii*) offers an attractive alternative for expressing viral CPs to obtain VLPs due to its ease of manipulation and high-level expression of recombinant proteins [61]. *P. pastoris* can achieve high cell growth in minimal media and maintain product stability in prolonged processes, making it suitable for developing bioreactor processes for biomedical relevant recombinant proteins [63]. Another well-defined and preferred expression system for producing chemicals and proteins is *Saccharomyces cerevisiae* [64]. Both yeasts offer the additional advantage of secretory expression of heterologous proteins, which is not typically available in the *E. coli* system [65].

Here, we demonstrate for the first time the self-assembly into VLPs of sobemovirus CP expressed in the methylotrophic yeast *P. pastoris*. Using RGMoV CP, we highlight the significant impact of the expression system on *in vivo* VLP formation. Our results show that VLPs formation can be achieved not only by switching the expression system from bacterial to yeast but also by transitioning from episomal to genome-integrated systems. Additionally, we demonstrated that the BacterioMatch II Two-Hybrid system is a valuable tool for studying nucleic acid-binding proteins. This allowed us to conclude that RGMoV CP has higher binding activity to ssDNA than RYMV or CfMV CPs. Leveraging this property, we encapsulated a type A CpG TLR9 agonist—G10—demonstrating the potential of RGMoV VLPs as nanocontainers or vaccine platforms with interior modification capabilities for enhanced immune response.

Moreover, our RNA-Seq data indicates that sobemovirus-derived VLPs predominantly package their CP mRNA, irrespective of the expression system used. This characteristic can be utilized for the functional encapsulation of nucleic acids.

## 2. Results and discussion

### 2.1. CPs expression analysis from episomal expression system in *E. coli* and *S. cerevisiae*

Since the first production and purification of VLPs in *E. coli* [66-69], the interest and need for novel VLPs have continued to grow. Plant viruses and their VLPs are valuable tools in biotechnology due to their well-characterized properties over several decades, including high yields, robustness, ease of purification, and the ability to modify surfaces both interiorly and exteriorly. These properties have attracted interest in medicine and nanotechnology for applications such as the delivery of anticancer compounds, targeted bioimaging, and vaccine production [70].

In this study, we aimed to characterize side-by-side three structurally well-described sobemoviruses by successfully expressing three soluble sobemovirus CPs – CfMV, RYMV[23] and RGMoV – in the *E. coli* expression system. All CPs were purified using a sucrose gradient. Analysis of the gradient fractions by SDS-PAGE and WB with polyclonal antibodies derived against native viruses or VLPs indicated CP signals in the sucrose fractions from 40% to 50%, consistent with our previous results for plant virus CP-derived VLP purifications [22, 23, 25, 36, 71, 72]. Fractions with CP signals were pooled for further purification and analysis. Purified CPs and a control sample (empty expression vector) were analyzed by TEM. As expected, the analysis revealed that CfMV and RYMV CPs self-assembled into VLPs, while RGMoV CP formed large aggregates (Fig. 1).

**Figure 1.**
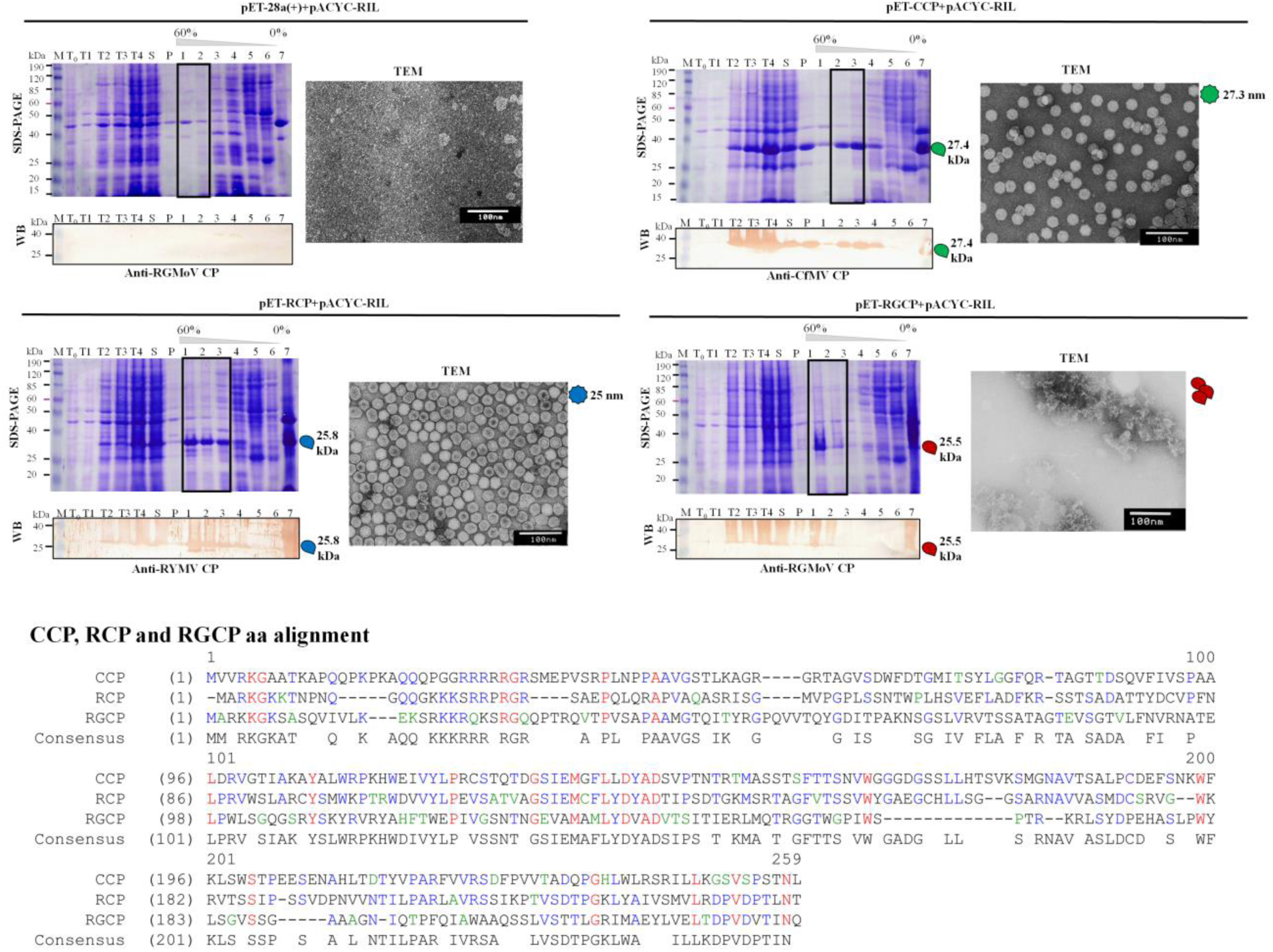
Expression and purification analysis of empty pET-28a vector, CCP, RCP and RGCP. T_0_ – total cell lysate before induction; T1 – total cell lysate after 2 h expression; T2 – total cell lysate after 4 h expression; T3 – total cell lysate after 6 h expression; T4 – total cell lysate after 20 h expression; S – supernatant after cell ultrasound sonication and sample clarification; P – pellet after cell ultrasound sonication and sample clarification; 1 – sucrose gradient fraction containing 60% sucrose; 2 – sucrose gradient fraction containing 50% sucrose; 3 – sucrose gradient fraction containing 40% sucrose; 4 – sucrose gradient fraction containing 30% sucrose; 5 – sucrose gradient fraction containing 20% sucrose; 6 – top fraction after sucrose gradient; 7 – pellet after sucrose gradient; M – pre-stained protein marker (BenchMark™ Pre-stained Protein Ladder, Thermo Fisher Scientific, USA). SDS-PAGE – sample analysis by 12.5% dodecyl sulfate–polyacrylamide gel electrophoresis and gel staining with Coomassie blue R250 stain; WB – Western blot (primary antibodies: rabbit anti-CfMV CP (1:1000), anti-RYMV CP (1:1000), anti-RGMoV CP (1:1000); secondary antibodies: horseradish peroxidase-conjugated anti-rabbit IgG (1:1000, Sigma-Aldrich, USA); TEM – transmission electron microscopy images with 1% uranyl acetate negative stain. SDS-PAGE tracks marked in the rectangle were used for further purification. CCP identity to RCP ∼ 30% [33], CCP identity to RGCP 12%, and RGCP identity to RCP ∼ 21% [34].

After unsuccessful VLP self-assembly in the *E. coli* expression system, we performed RGCP expression in two *S. cerevisiae* strains, AH22 and DC5. Since the yeast expression system can result in glycosylation of the target recombinant protein and, according to N-glycosylation analysis by NetNGlyc - 1.0 [73], RGCP Asp^94^ is predicted to be N-glycosylated (Fig. S1). Therefore, using *S. cerevisiae* as an expression system we e decided to evaluate the possible glycosylation effect on CP in VLP self-assembly.

Expression vector pFX7 designed for *S. cerevisiae* [74] has been successfully used for several RNA bacteriophage CPs expression [75-77]. *S. cerevisiae* has been used as an expression system for several plant virus CPs for VLPs purification [78, 79]. *S. cerevisiae*, due to its fast-growing, robust nature and well-known genetics, is one of the widely used single-cell eukaryotic model systems for heterologous protein expression for various purposes – vaccines, chemicals, hormones, and biofuels [80]. RGCP expression was tested in YPD media with formaldehyde.

Both strain cell total lysates were analyzed in WB with polyclonal antibodies against RGMoV, revealing RGCP expression but at a lower level than in the *E. coli* expression system (Fig. S1). Yeast cells were disrupted by French press in 15 mM KH_2_PO_4_ buffer, and the soluble protein fraction was purified by sucrose gradient. After fraction analysis by WB, the RGCP specific signal was detected in two sucrose gradient fractions – 50% and 40% (Fig. S1), similar to the *E. coli* case (Fig. 1). Both sucrose fractions containing RGCP signal were pooled and concentrated by ultracentrifugation. TEM analysis of the sample revealed icosahedral particles approximately 40 nm in diameter with an empty core (Fig. S1), which is unusual compared to the previously reported 28 nm for native RGMoV virions and 25 nm from a 3D structure model prediction [34, 81, 82]. Since virus CP-derived VLPs frequently encapsulate the mRNA of the *CP* [35], it can be used as a marker to identify the origin of VLPs. To determine the origin of the isolated viral particles, RAP-PCR was performed. Sanger sequencing analysis revealed that the icosahedral, ∼40 nm particles observed in TEM contained sequences from the *S. cerevisiae* helper virus L-A [83], rather than *RGCP* mRNA. This suggests that RGCP expression may have provoked helper virus replication, as changes in the L-A virus level in yeast cells occur under different metabolic conditions [84-86]. L-A virus is common in a various wild, industrial and laboratory yeasts [87-89]. Alternatively, the RGCP expression level might have been too low for effective VLP assembly. Consequently, switching the expression host to *S. cerevisiae* did not result in the successful formation of VLPs from RGCP.

### 2.2. RGMoV CP expression in *P. pastoris*

The third approach to test RGCP VLP formation involved switching to a genome-integrated expression system using the methylotrophic yeast *P. pastoris* [90]. This expression system is well-established for producing VLPs from various sources, including bacteriophages, plant viruses and the hepatitis B virus core protein (HBc) [61, 75, 76, 91]. The *P. pastoris* expression system has been used for over 30 years due to its easy genetic manipulation, high-frequency DNA transformation, functional complementation cloning, and the ability to produce high levels of intra- and extracellular proteins. Moreover, it can perform higher eukaryotic protein modifications such as glycosylation, disulfide bond formation, and proteolytic processing, making it invaluable for both research and industrial production of recombinant proteins [92].

To explore the potential of *P. pastoris* for the expression of RGMoV CP, we performed electroporation to introduce the plasmid construct containing *CP* gene into *P. pastoris* cells for genome integration. Post-transformation, *P. pastoris* cells were initially selected on His-medium. For further selection, cells were plated on YPD agar with increasing concentrations of gentamicin (G418; 2 mg/ml and 4 mg/ml). Six colonies that grew on 4 mg/ml G418were picked for subsequent expression studies as clones with a G418 resistance level were found at 2–3% frequency, suggesting insertions with increased copy number [76]. Six clones were selected for RGMoV CP expression analysis. Protein extracts from these clones were subjected to 12.5% SDS-PAGE followed by Coomassie staining and WB using polyclonal anti-RGMoV CP antibodies (Fig. 2). Additionally, PCR was performed using genomic DNA isolated from selected *P. pastoris* clones to confirm the presence of the *CP* gene (the quantity of the inserted *CP* gene copies was not calculated). Four out of the six clones demonstrated CP expression visible by Coomassie staining (Fig. 2). All six clones expressed the RGMoV CP, though the expression levels varied, with some clones showing lower expression levels not detectable by Coomassie staining (Fig. 2). PCR analysis of genomic DNA demonstrated the presence of the *CP* gene in all six clones (Fig. 2), consistent with the WB results (Fig. 2). These findings suggest that the expression of RGMoV CP in *P. pastoris* is dependent on the copy number of the integrated gene, and the clones with higher gene multiplicity exhibit stronger expression levels.

**Figure 2.**
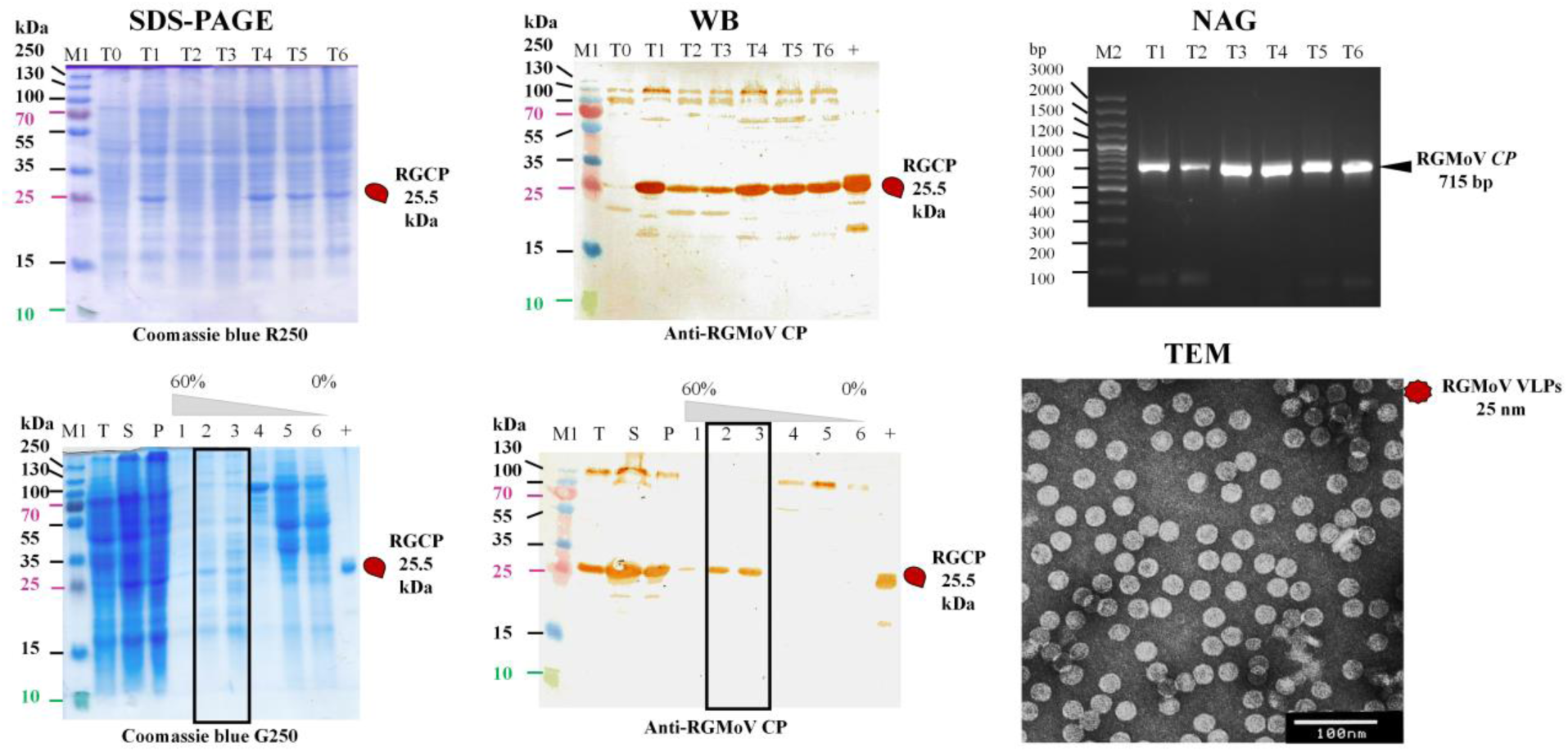
RGCP expression and purification analysis (*P. pastoris* system). SDS-PAGE – 12.5% dodecyl sulfate–polyacrylamide gel electrophoresis and gel staining with Coomassie blue R250 or G250 stain; WB – Western blot analysis (primary antibodies: anti-RGMoV CP (1:1000); secondary antibodies: horseradish peroxidase-conjugated anti-rabbit IgG (1:1000, Sigma-Aldrich, USA); NAG – 0.8% native agarose gel stained with ethidium bromide for PCR analysis of *P. pastoris* expression clones for RGMoV *CP* gene sequence in chromosomal DNA (amplicon length 715 bp); TEM – transmission electron microscopy images with 1% uranyl acetate negative stain. SDS-PAGE tracks marked in the rectangle were used for further purification. M1 – PageRuler™ Prestained Protein Ladder, 10-250 kDa (Thermo Fisher Scientific, USA); M2 – GeneRuler 100 bp Plus DNA Ladder (Thermo Fisher Scientific, USA); T_0_ – total cell lysate before induction; T1-6 – total cell lysate of expression clones after expression; T – total cell lysate after disruption by Covaris; S – supernatant after cell ultrasound sonication and sample clarification; P – pellet after cell ultrasound sonication and sample clarification; 1 – sucrose gradient fraction containing 60% sucrose; 2 – sucrose gradient fraction containing 50% sucrose; 3 - sucrose gradient fraction containing 40% sucrose; 4 – sucrose gradient fraction containing 30% sucrose; 5 – sucrose gradient fraction containing 20% sucrose; 6 – sample fraction after sucrose gradient; “+” – native RGMoV as positive control.

This aligns with the general understanding that higher gene dosage can lead to increased protein production in heterologous systems. The use of *P. pastoris* for the expression of RGMoV CP proved to be promising. The selection of clones on high G418 concentrations allowed for the identification of strains with multiple copies of the *CP* gene, resulting in varying expression levels.

Yeast biomass from expression clones No. 1, 4, 5, and 6 were pooled and disrupted using Covaris. The soluble protein fraction, after clarification by low-speed centrifugation, was purified using a sucrose gradient. All fractions were analyzed by 12.5% SDS-PAGE and WB. The analysis revealed that the CP was located in the 50% and 40% sucrose gradient fractions (Fig. 2), similar to the patterns observed in *E. coli* (Fig. 1) and *S. cerevisiae* (Fig. S1). The sucrose gradient fractions containing the CP signal were pooled, dialyzed, and concentrated by ultracentrifugation. The resulting pellet was solubilized and measured on a NanoDrop-1000, and samples were analyzed in 0.8% NAG. The NanoDrop-1000 measurements showed a maximum peak at 260 nm, similar to the CCP and RCP cases [23]. NAG analysis also revealed a distinct NA band that did not disappear after treatment with Benzonase or DNase, and RNase T1/A (Fig. S2), indicating a protein-NA complex. NAG staining with G250 showed a protein signal overlapping with the NA signal detected by ethidium bromide. Only after DNase and RNase T1/A treatment did some of the protein signal migrate differently from bensonase and nontreated samples in NAG (Fig. S1), likely due to VLP disassembly caused by RNA degradation by RNase. Native RGMoV virions after treatment with RNases and subsequent RNA isolation demonstrated low RNA content not suitable for NGS library preparations [81]. RNase treatment approach had been used to obtain empty VLPs for heterologous cargo encapsulation by diffusion through holes of ∼2 nm diameter present in the capsid structure [93] or for the impact investigation of encapsulated RNA to immune system response [16, 94, 95].

TEM analysis confirmed that RGCP self-assembled into VLPs, which did not differ visually from purified native virions (Fig. 3). DLS analysis demonstrate homogenious particle population with a hydrodynamic diameter 28.51 nm. whereas parallel analysis of native RGMoV particles was found 26.63 nm). It corresponds well with reported 28 nm diameter of native RGMoV particles [82] and a 3D structure model predicting a 25 nm diameter [34]. MALDI TOF mass spectrometry analysis of purified RGMoV virion and VLPs revealed that both variants lack the first methionine (Met), with the virion-forming CP mass at 25.466 kDa and VLP-forming CP at 25.433 kDa (Fig. 3). This aligns with the calculated molecular mass without Met (25.49 kDa) and the mass with the first Met (25.6 kDa) [96]. A similar observation was made for Qβ CP expression in *E. coli*, *S. cerevisiae*, and *P. pastoris* systems [76], as well as for CfMV, RYMV CPs [23], and RGMoV protease [97] expressed in *E. coli*. This occurs because the next aa after Met is a small AA residue [98, 99]. As the detected molecular mass of RGMoV CP was close to predicted, that demonstrates the RGMoV CP for maturations and proper folding posttranslational modifications are not required despite predicted glycosylation of Asp^94^ (Fig. S1) As a next we tested a thermal stability of sobemovirus VLPs using the fluorescent dye Sypro-Orange [100] and a real-time PCR device with a DNA melting point determination program. Previous research has shown that this method effectively characterizes the termal stability of purified VLPs and shows structural changes depending on the temperature [25, 36]. The results indicated that the overall stability of the VLPs was not significantly affected. The fluorescence maximum for RGMoV VLPs in standard buffer (15 mM KHPO_4_ pH 5.5) differed by only 4 °C compared to the native virus (Table 1; Fig. 4). A decrease in stability was observed only in samples with added EDTA, where the melting temperature decreased by at least 10 °C (Table 1; Fig. 4). EDTA, a chelating agent, forms very stable complexes with most divalent and trivalent metal ions [101]. Divalent cations play a significant role in the stability of sobemovirus virion [102, 103].

**Figure 3.**
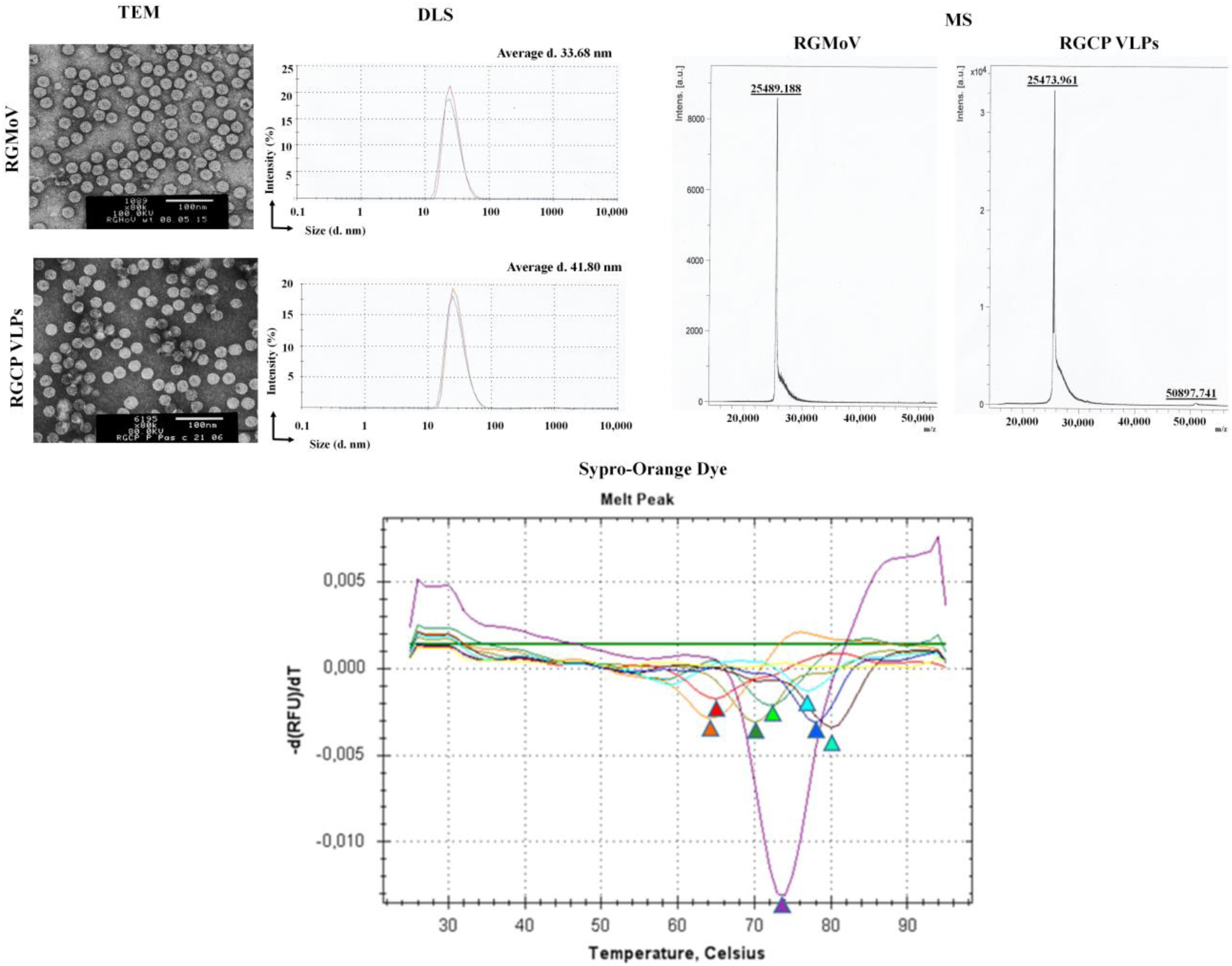
Native RGMoV and *P. pastoris* generated RGCP VLP analysis. TEM – transmission electron microscopy analysis by negative stain with 1 % uranyl acetate; DLS – dynamic light scattering analysis of hydrodynamic diameter; MS – mass spectrometry analysis of CPs; Sypro-Orange Dye – analysis of CPs melting curve on RT-PCR device with the fluorescent dye Sypro-Orange (Sigma-Aldrich, USA); 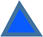– RGMoV in standart buffer (15 mM KHPO_4_ pH 5.5); 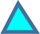 – RGMoV in standart buffer supplemented with 50 mM DTT; 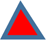 – RGMoV in standart buffer supplemented with 25 mM EDTA; 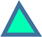 – RGMoV in standart buffer supplemented with 0.5 M NaCl; 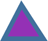 – RGCP VLPs in in standart buffer (15 mM KHPO_4_ pH 5.5); 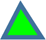 – RGCP VLPs in in standart buffer supplemented with 50 mM DTT; 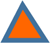 – RGCP VLPs in in standart buffer supplemented with 25 mM EDTA; 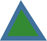 – RGCP VLPs in in standart buffer supplemented with 0.5 M NaCl.

**Figure 4.**
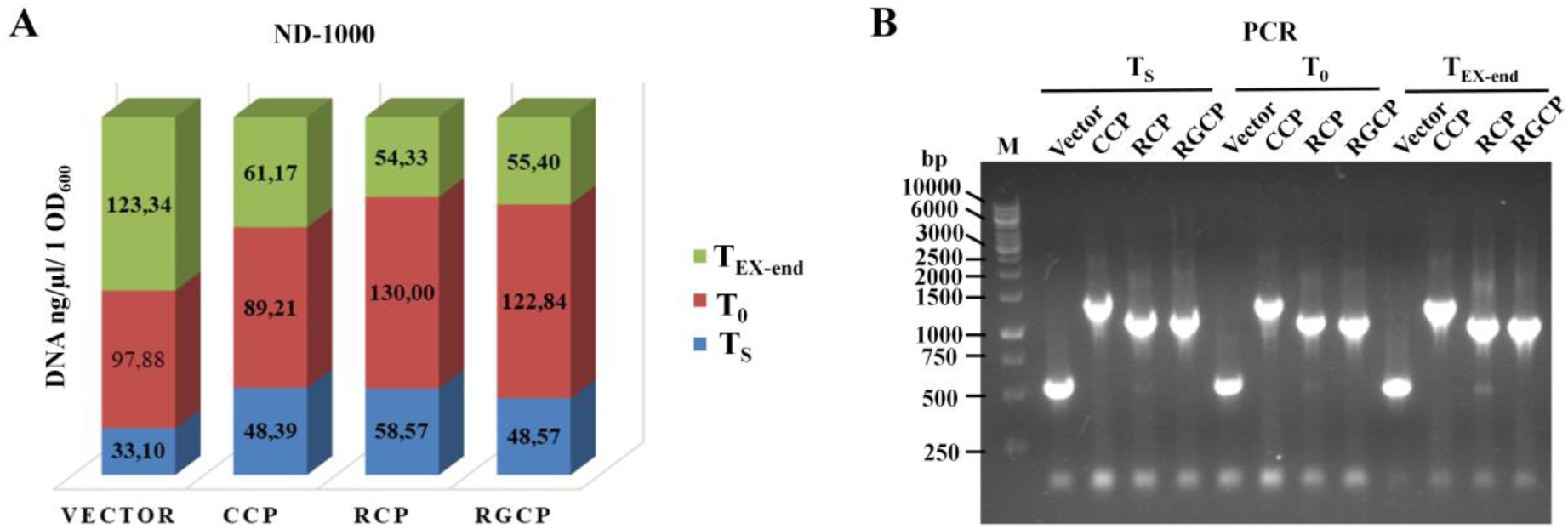
Plasmid and PCR analysis during the CPs expression experiment. T_S_ – plasmid DNA isolated from seed culture; T_0_ – plasmid DNA isolated from cell culture at OD=0.8 directly before induction with 0.2 mM IPTG; T_EX-end_ – plasmid DNA isolated from cells after 20 h expression, the end of expression process; M – DNA 1 kb size marker (Thermo Fisher Scientific, USA). A – graphical overview of purified plasmid concentrations in three time points of expression experiment calculated on 1 OD_600_; B –PCR product analysis in 0.8% NAG stained with ethidium bromide.

**Table 1.**
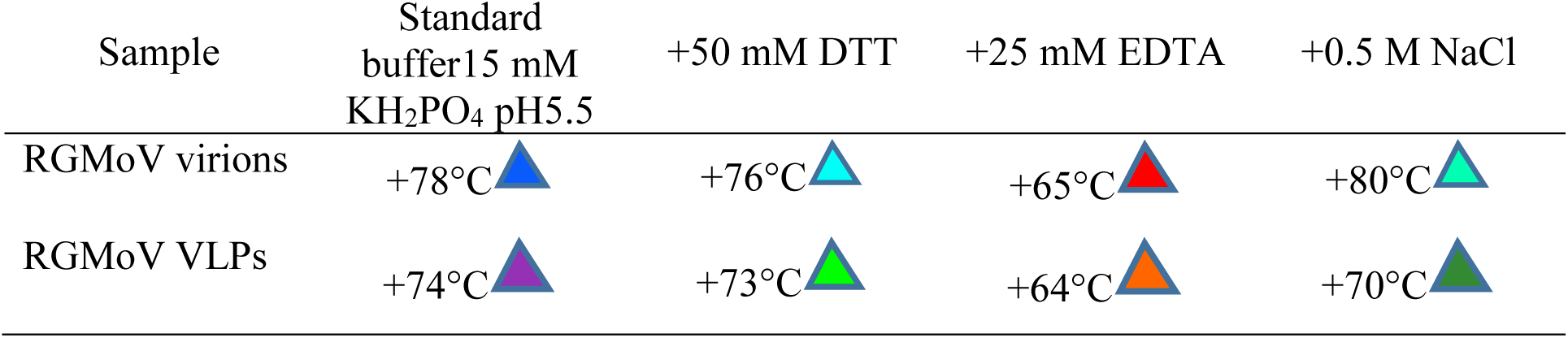
Analysis of RGMoV virion and VLPs thermal stability.

| Sample | Standard<br>buffer15 mM<br>KH<sub>2</sub>PO<sub>4</sub> pH5.5 | +50 mM DTT | +25 mM EDTA | +0.5 M NaCl |
| --- | --- | --- | --- | --- |
| RGMoV virions | +78°C | +76°C | +65°C | +80°C |
| RGMoV VLPs | +74°C | +73°C | +64°C | +70°C |

### 2.3. Characterization of factors influencing the VLP self-assembly

Published research typically focuses on success stories, as unsuccessful attempts raise many questions and are harder to publish. Nevertheless, we decided to perform a deeper analysis using RGMoV CP as a model to investigate why we were unable to obtain self-assembled VLPs from RGCP when expressing it in *E. coli* or *S. cerevisiae* systems. Our experiments included several steps. First, we monitored the expression step, comparing empty expression vectors, CCP, RCP, and RGCP OD_600_ changes during expression to assess potential toxicity (1), plasmid stability tests to evaluation of plasmid stability at the start of the culture and the end of expression (2), and monitoring the presence of CP and promoter sequences in plasmids extracted at different expression time points (3). The second part involved purified CP analysis divided into two parts based on the first part’s results: protected NA tests for detection of encapsulated nucleic acids using PCR or RT-PCR (4) and CP gel-shift analysis for assessment of CP binding to ssDNA and dsDNA (5). The third part involved CP analysis in the BacterioMatch II Two-Hybrid system for investigation of CP-NA binding interactions *in vivo* (6).

It is known that delays in cell growth can be associated with toxicity of expressed proteins [52]. To evaluate possible toxic effect of RGCP’s on cells and its impact on CP expression and VLP formation, we compared CPs and empty expression vectors by monitoring cell OD_600_ at different growth stages—during the first four hours after inoculation of the start culture (T_0_), during a six-hour period after induction with IPTG, and at the end of expression (T_EX_). OD_600_ monitoring revealed no significant delay in cell growth for CPs (Fig. S3).

Plasmid stability tests were performed for the seed culture used for inoculation (T_S_) and the expression end culture (T_EX-end_). The number of living cells was determined using basic microbiological techniques, calculated to 1 OD unit. Results showed no decrease in colony number at the start point, with similar cell growth on agarized LB media without or with both antibiotics from T_S_ culture. However, interesting results were observed when analyzing plates from T_EX-end_. In the CCP case, there was no significant reduction in plasmid-containing cells, demonstrating low toxicity and stability of the expression vector, correlating with high CCP expression levels. Dramatically opposite results were observed for RCP and RGCP, with nearly 100% loss of the expression vector in T_EX-end_ samples when comparing plates with Km, Km/CAM and without antibiotics. In the RCP case, 12.42% of Km-resistant cells survived in the T_EX-end_ sample, but only 0.008% survived in the RGCP case (Table 2). This could suggest a significant impact of RCP and RGCP toxicity on cells, leading to plasmid instability, especially for RGCP, which could explain the lack of VLP formation, though similar RCP conditions did not inhibit VLP self-assembly or expression level.

**Table 2.**
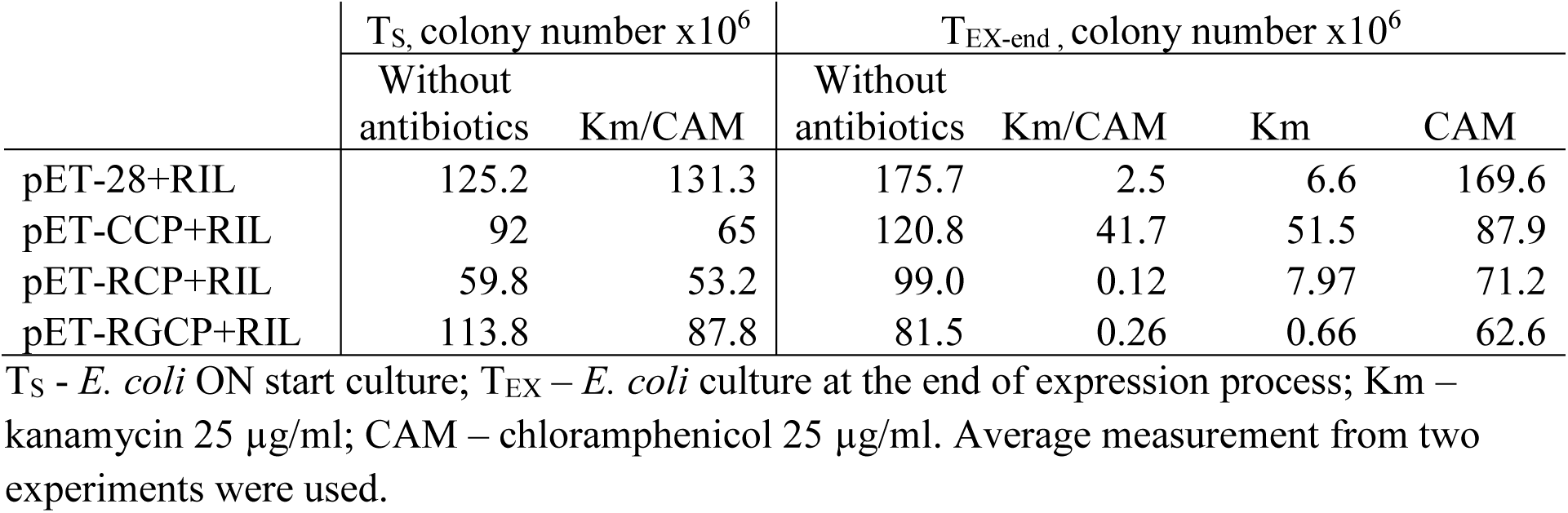
Measurement of survived bacteria on 1 OD unit during CP expression.

Due to the instability of RGCP and RCP expression plasmids, which may be caused by BL21(DE3) cells shutting off the pET system via mutations or by down-regulating expression due to protein toxicity that accumulates over long induction times (20 h) [104-106], we performed additional tests to evaluate the possible changes in the plasmid *T7* promoter and *CP* gene regions by PCR for plasmids extracted at different expression time points. Purified NA concentrations did not show critical changes during the experiment. Some minor differences were observed for RGCP, where NA at T_0_ was higher than in CCP and empty expression vector cases but lower after T_EX_ (Fig. 4A). We performed PCR with primers located in the expression vector—direct primer 220 nt upstream of the promoter (pET-220; Table 3) and reverse primer 80 nt after the *CP* stop codon (pET-R; Table 3) creating PCR amplicons of 549 bp (empty vector), 1277 bp (vector with CCP), 1147 bp (vector with RCP), and 1147 bp (vector with RGCP). The obtained PCR amplicons did not differ in length during the experiment (Fig. 4B).

**Table 3.**
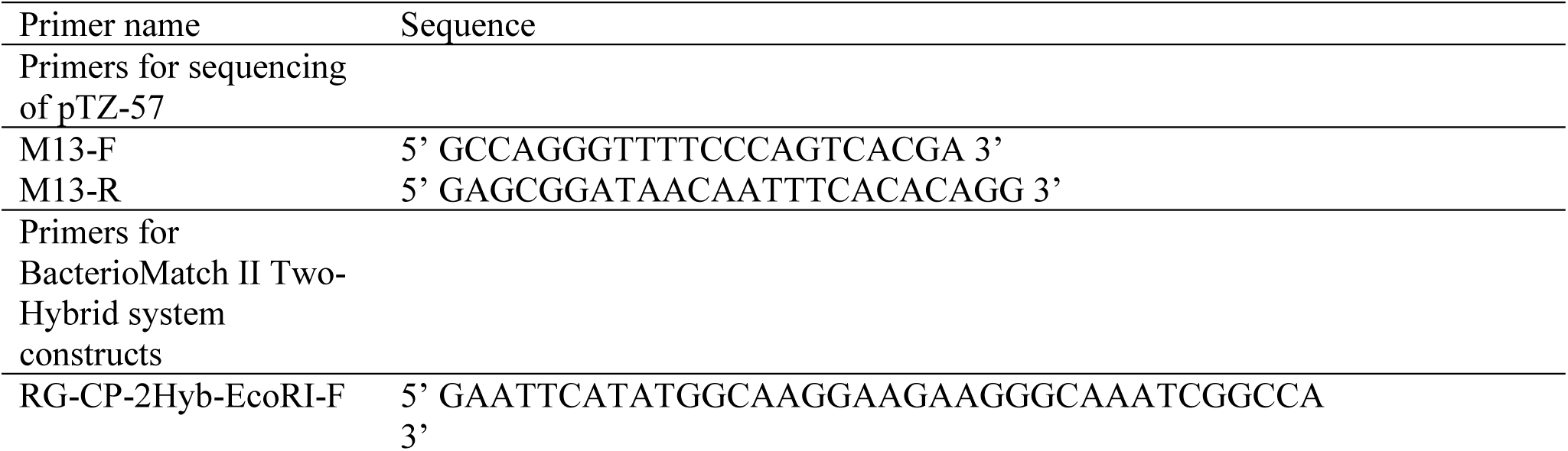
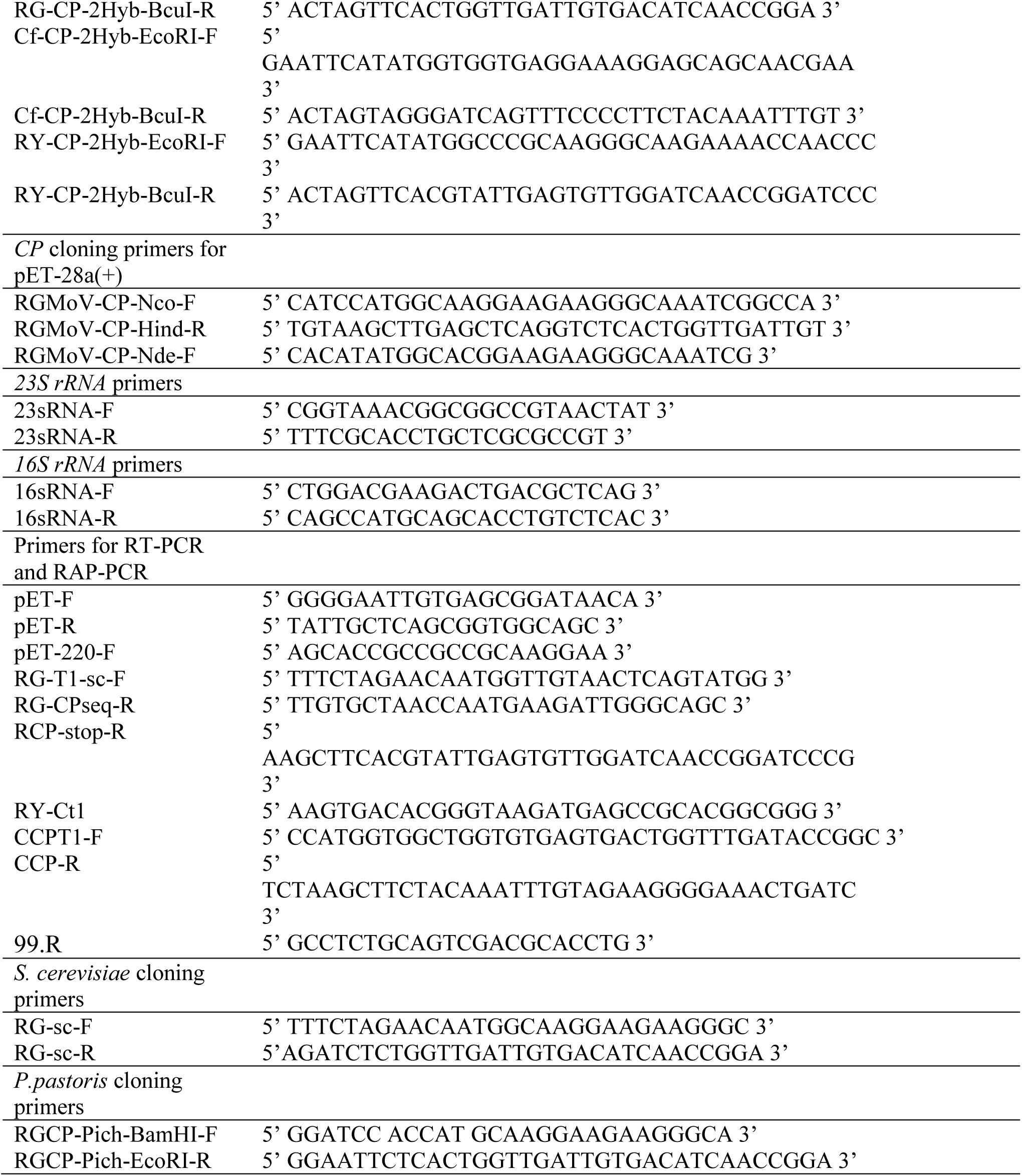
Primers used in this work.

All PCR products from pET-CCP, pET-RCP, and pET-RGCP were cloned into pTZ-57R/T for Sanger sequencing. Sequencing data did not reveal expression-influencing changes in the promoter region or *CP* sequence, demonstrating that proposed hypothesis of plasmid instability due to long toxic effect of CPs is not the reason that had led to misassemble of RGCP VLPs. The observed low maintenance of pET28a(+) plasmid during plasmid stability test (Table 2) in vector, RCP and RGCP 20 h expression samples is mostly due to pET plasmid itself. The efficiency of plasmid maintenance depends on several factors, but increased plasmid maintenance did not result in higher production titers [107, 108].

Sobemovirus CPs’ N-terminal part contains a R domain rich in arginine, lysine, proline, and glutamine, essential for CP-RNA contacts [109, 110]. The total number of positive charges associated with the RNA is around 2,340, sufficient to cancel about half of the negative charges of the NA. SCPMV R domain’s first 54 aa expressed in *E. coli* showed nonspecific RNA binding activity *in vitro* [26]. SeMV CP N-terminally truncated version by 22 aa assembled into stable *T = 3* VLPs resembling native SeMV, indicating the dispensability of the N-terminal 22 aa for *T = 3* assembly. A 36 aa N-terminally truncated SeMV CP version assembled into *T = 1* and pseudo *T = 2* VLPs, while a 65 aa N-terminally truncated version formed *T = 1* VLPs [35]. In RGMoV, removing the N-terminal 54 aa did not result in VLP formation (data not shown). All SeMV N-terminally truncated variants encapsidated *23S rRNA* and *CP* mRNA, suggesting additional NA binding sites may be involved in RNA-CP interactions [35]. CfMV CP (native and *E. coli* expressed) NA studies *in vitro* demonstrated that both bound ssRNA nonspecifically and were selective for ssRNA over dsDNA molecules [111]. Therefore, we tested purified CPs for encapsulated or CP-protected NA to identify possible differences in VLPs packaged NA composition versus CP aggregates. SeMV VLPs have been reported to package *CP* mRNA and *23S rRNA* transcripts [35]. For this experiment, we used previously purified samples, including an empty expression vector as a negative control. Prior to NA isolation, we treated purified samples with benzonase to eliminate unprotected NA that could influence results. We first tested purified material for possible plasmid DNA presence by direct PCR with plasmid-specific primers (pET-F and pET-R (vector – 295 bp, CCP – 1023 bp, RCP – 893 bp; RGCP – 893 bp; Table 3). PCR results showed only the RGCP sample was positive for plasmid DNA (Fig. 5).

**Figure 5.**
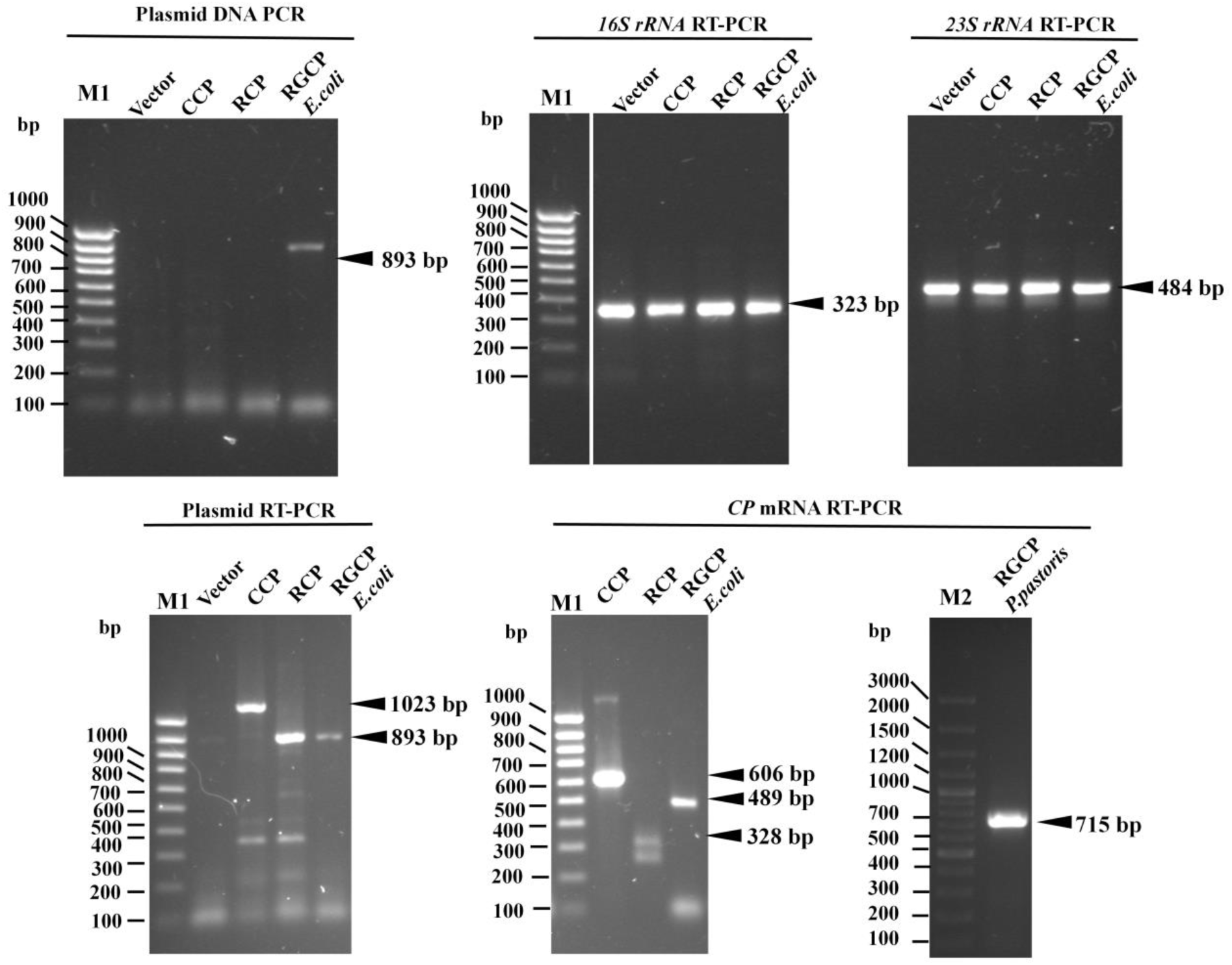
Plasmid DNA and *16S, 23S rRNA* and *CP* transcript detection by PCR or RT-PCR in isolated nucleic acid pool from expression experiment samples. M1 – GeneRuler 100 bp DNA Ladder (Thermo Fisher Scientific, USA); M2 – GeneRuler 100 bp Plus DNA Ladder (Thermo Fisher Scientific, USA).

This could explain the absence of VLP formation due to plasmid dsDNA size (5966 bp), or partial ssDNA form due to replication peculiarities [112]. Additionally, we transformed all purified samples into XL1 Blue cells to detect possible replicating plasmids. No colonies were detected on LB plates, indicating that the binding RGCP to the plasmid DNA which can be a partial ssDNA form of the expression plasmid. This explains the observed low number of plasmid copies after expression instead of the instability of the plasmid itself as initially proposed (Fig. 5).

RT-PCR was used to detect *CPs* mRNA, *16S*, and *23S rRNA* transcripts. cDNA was obtained by random hexamer primers followed by PCR with transcript sequence-specific primers (*CCP* – CCPT1-F and CCP-R (606 bp), *RCP* – RCP-stop-R and RY-Ct1 (328 bp), *RGCP* – RG-T1-sc-F and RG-CP-seq-R (489 bp) for *E. coli* purified aggregates or RGCP-Pich-BamHI-F and RGCP-Pich-EcoRI-R for VLPs purified form *P. pastoris* (715 bp), *16S rRNA* – 16sRNA-F and 16sRNA-R (323 bp), *23S rRNA* - 23sRNA-F and 23sRNA-2470R (484 bp); Table 3). For RGCP VLPs, only *CP* mRNA transcript detection was performed. RT-PCR analysis in 0.8% NAG showed corresponding PCR amplicon for *16S* and *23S rRNA* in all samples purified from *E. coli* (Fig. 5). This could be due to ribosome size, which corresponds to VLP size and sedimentation properties in sucrose gradients similar to VLPs [113], leading to their co-purification. The rRNA is protected by ribosomal proteins from benzonase activity, similar to VLPs and CP aggregates. EDTA can disrupt ribosome complexes, but is not preferred due to its effect on sobemovirus VLP and native virus stability [102]. Lokesh and coauthors [35] proposed that the detected *23S rRNA* in SeMV VLPs RNA samples might be due to ribosomes co-purified with VLPs in sucrose gradients [113]. *CP* mRNA corresponding gene-specific transcript PCR product was detected in all CP samples regardless of the expression host (Fig. 5). To avoid biases inherent in sequence-specific PCR, we used HTS for analyzing encapsulated RNA profiles. This approach allowed us to identify nucleic acid sequences without the need for sequence-specific primers, offering a more comprehensive and semiquantitative analysis of RNA sequences [114]. RNA-Seq data were analyzed to determine the proportion of reads corresponding to *CP* mRNAs and host RNA transcripts, providing insights into the specificity and preference of viral nucleic acid sequences. The RNA-Seq data revealed that the majority of mapped reads corresponded to CP mRNA, with CCP at 67%, RCP at 38%, and RGCP at 100% (Fig. 6). This demonstrated a strong preference and specificity for viral nucleic acid sequences regarding expression host RNA transcripts. This demonstrated the viral NA sequence preferability and specificity regarding expression host RNA transcripts. It could be explained due to fact that viral NA possess e structure density [115]. For CCP, the viral mRNA constituted 67% of the mapped reads. Host origin transcripts formed 14% of the unique mapped reads with coverage of 1. The main host transcript was *lacI* at 1%, while 13% of host transcripts represented 8553 different genes. Plasmid transcripts were abundant, with *CamR* at 8%, *KanR* at 6%, and *lacI* at 3% (Fig. 6). In the case of RCP, viral mRNA accounted for 38% of the mapped reads. Host origin transcripts were more abundant, forming 37% of unique mapped reads with coverage of 1. The most abundant host transcripts were *lacI* and *sslA*, each representing 1% of identified transcripts. Plasmid transcripts were also prevalent, with *KamR* at 11% and *lacI* at 12% of mapped reads (Fig. 6). Notably, neither CCP nor RCP transcripts included *16S rRNA* or *23S rRNA*. For RGCP, almost all mapped reads, except for two host-derived sequences, were entirely from *CP* mRNA, showing a high specificity for viral RNA (Fig. 6).

**Figure 6.**
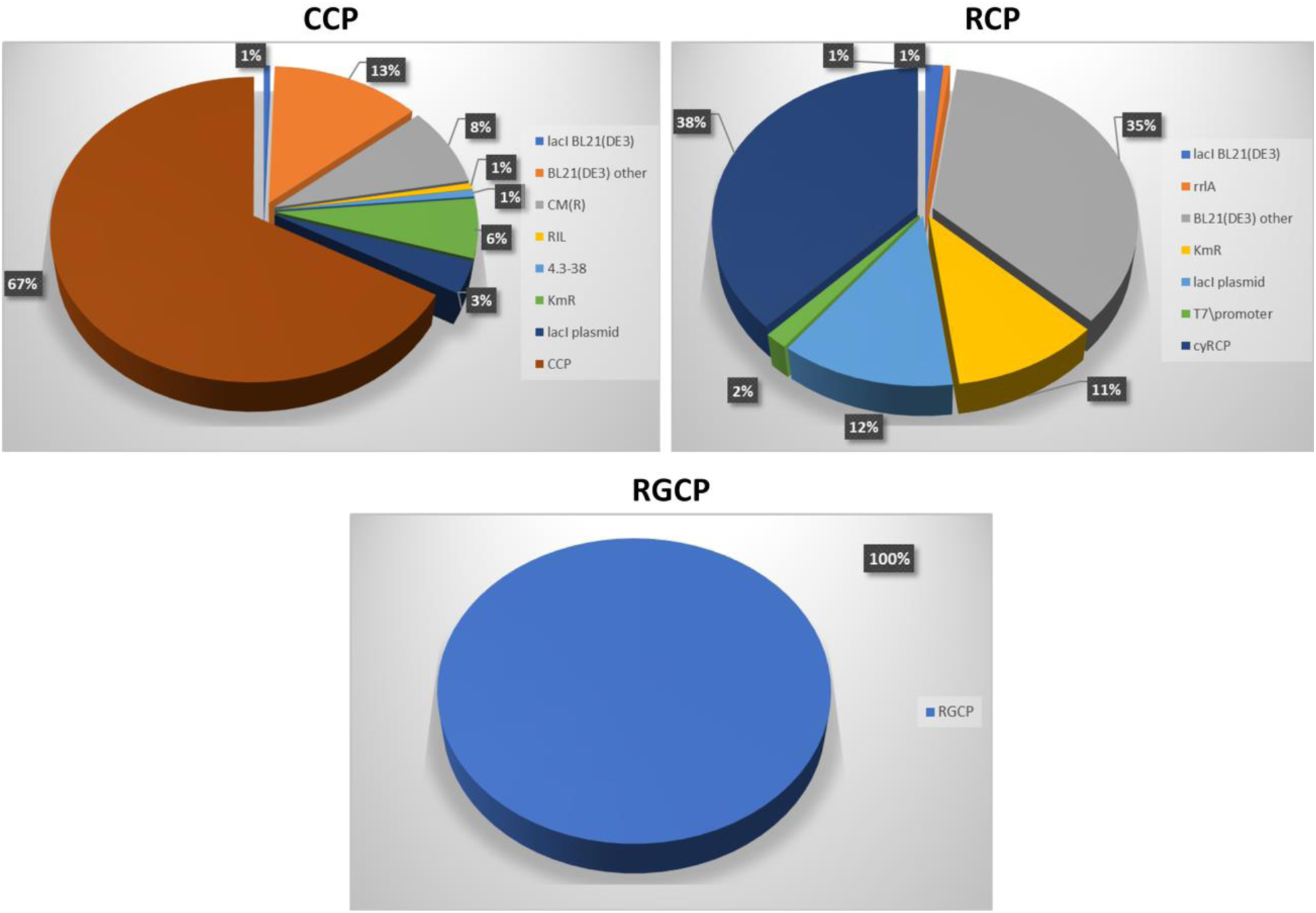
The analysis of mapped RNA transcripts encapsulated in CCP, RCP and RGCP VLPs.

The RNA-Seq data demonstrated a clear preference and specificity for viral nucleic acid sequences, particularly in the case of RGCP, where the viral mRNA predominated almost entirely. CCP and RCP samples showed a substantial proportion of host-derived transcripts, indicating a lower specificity compared to RGCP. The presence of host RNA in CCP and RCP samples suggests that viral CPs encapsulate a mixture of viral and host RNAs, potentially influenced by the structural density of viral nucleic acids. These findings underscore the capability of RNA-Seq to provide a detailed profile of encapsulated RNA, revealing both the specificity of viral CPs for their mRNAs and the extent of host RNA inclusion in the assembled particles. The predominance of viral RNA in RGCP indicates a high specificity, which could be beneficial for future applications in VLP-based delivery systems.

A similar situation was observed for HBc VLPs derived from the *E. coli* expression system, where the majority of RNA transcripts were from the HBc coding sequence, but *23S* and *16S* rRNA were detected in low abundance [116]. Similar observations were made for other VLPs as well [95, 116].

Given the observed RGCP binding activity to plasmid DNA, we tested all three sobemovirus CPs (RGMoV, CfMV, and RYMV) for NA binding using the BacterioMatch II Two-Hybrid system. This system’s known limitation—its propensity to detect NA binding proteins as interactors with the empty bait plasmid—was leveraged to model NA binding *in vivo*. *CP* coding sequences were cloned into the target plasmid, with the empty bait plasmid serving as the NA binding test. *E. coli* cells expressing a CP that binds the bait plasmid DNA will grow, while those that do not bind will not grow. The BacterioMatch II Two-Hybrid system results showed that RGCP had a stronger binding ability to the bait plasmid compared to CCP and RCP (Fig. S4). Building on the BacterioMatch II Two-Hybrid system data and NA PCR experiment observations, we performed an additional test to identify whether RGMoV CP preferentially binds dsDNA or ssDNA. This was done using a gel shift assay in 0.6% NAG, with -N6H-RGCP purified by IMAC under denaturing conditions and subsequently refolded. Linearized pET-N6H-RGCP expression plasmid with HindIII or double digested with KpnI and BglII, and the PCR product of the RGMoV *CP* gene served as sources of dsDNA and ssDNA, respectively. The gel shift analysis revealed that RGMoV CP had a higher binding capacity to ssDNA than dsDNA (Fig. 7). None of the double-digested plasmid fragments showed a higher binding affinity to N6H-RGCP, suggesting that the interaction might be sequence-unspecific. These tests confirmed our hypothesis that RGMoV CP binds to plasmid DNA, especially ssDNA, which occurs during plasmid replication. This strong binding to plasmid ssDNA could led to RGCP binding to plasmid replication intermediates. The intermediates can be too large to be encapsulated in to VLPs. As a result, we have aggregates instead of VLPs. As shown above, expression system of *P. pastoris* system with a genome-integrated *CP* gene allows the VLP formation. In this system RGMoV CP obviously cannot bind DNA molecules.

**Figure 7.**
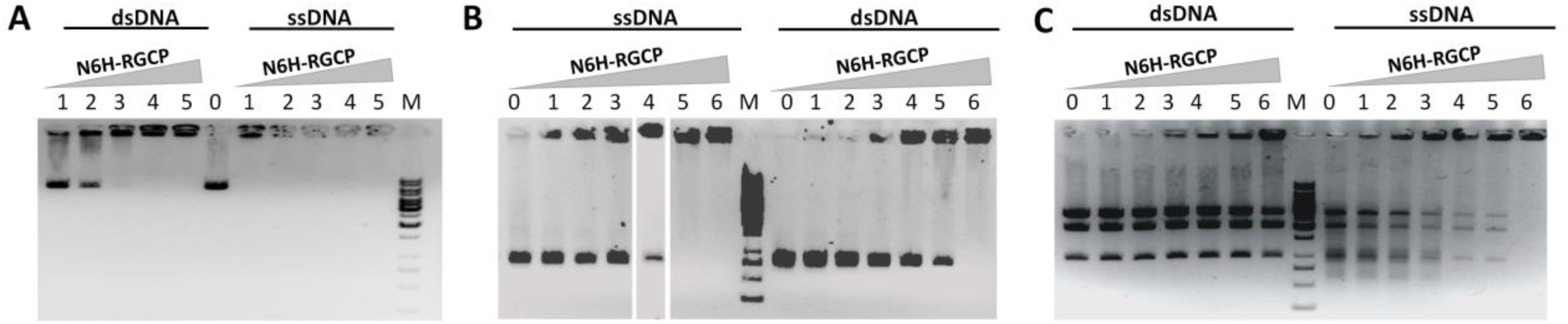
RGMoV CP binding test to dsDNA and ssDNA by mobility gel shift assay. A – N6H-RGCP with linearized pET-N6H-RGCP expression plasmid with HindIII (250 ng); B – RGMoV *CP* gene PCR product (250 ng); C – linearized pET-N6H-RGCP expression plasmid with KpnI and BglII (3.4 µg); 0 – plain vector DNA; 1 – N6H-RGMoV at 175 µg concentration; 2 – N6H-RGMoV at 250 µg; 3 – N6H-RGMoV at 500 µg concentration; 4 – N6H-RGMoV at 1000 µg; 5 – N6H-RGMoV at 2000 µg; 6 – N6H-RGMoV at 4000 µg; M – DNA 1 kb size marker (Thermo Fisher Scientific, USA).

The demonstrated RGCP binding activity to ssDNA may reflect a broader role for sobemovirus CPs in host replication or transcription suppression. CfMV CP contains two nuclear localization signals (NLS), indicating RNA transport to the nucleus [117]. RYMV CP’s first 22 aa also contain a sequence similar to NLS [118]. CfMV CP acts as an RNA silencing suppressor [119]. Post-transcriptional gene silencing (PTGS), an RNA-targeted homology-dependent RNA degradation process occurring exclusively in the cytoplasm, is a plant defense mechanism against viruses [120]. The observed interaction of RGMoV CP and other sobemovirus CPs with ssDNA, combined with NLS targeting the nucleus, suggests a possible role in PTGS suppression. Further *in planta* experiments are required to confirm this hypothesis.

The development of safe, effective, and immunogenic vaccines targeting various antigens necessitates appropriate adjuvants capable of modulating the immune response towards the desired outcome [121]. TLR ligands have shown promise as adjuvants for vaccines due to their safety and remarkable ability to enhance immunogenicity, with efficacy already proven in clinical settings [122, 123]. Depending on the NA encapsulated into VLPs, immunoglobulin G (IgG) isotype switching can be achieved [124]. TLR 3, TLR7/8, and TLR9 ligands, such as dsRNA, ssRNA, and CpG, especially in combination with delivery systems like nanoparticles and VLPs, have shown effective induction of cytotoxic T cell response, IgG class switching, and neutralizing antibody production for different pre-clinical vaccine prototypes targeting various indications [16, 93, 121, 125].

To explore the potential of RGMoV CP-derived VLPs as a vaccine platform that can be loaded with an additional adjuvant to target TLR9, we utilized the RGCP’s ability to bind DNA for nucleic acid encapsulation tests. After VLP disassembly, we performed reassembly with a type A CpG TLR9 agonist—G10 at three concentrations (3 µg/µl, 1.5 µg/µl, 0.75 µg/µl) via overnight dialysis. Sample analysis using 12.5% SDS-PAGE and Western Blot after ultracentrifugation showed that the pellet after ultracentrifugation and solubilization contained RGCP. Analysis using 0.8% NAG showed a nucleic acid signal captured by ethidium bromide and Coomassie blue R250 dye (Fig. 8). Further sample analysis by TEM demonstrated that 0.75 µg/µl G10 or no G10 after the reassembly process resulted in CP aggregates (Fig. 8). Only VLP reassembly was achieved at 3 µg/µl and 1.5 µg/µl G10 concentrations (Fig. 8). In samples reasembled after ultracentrifugation where G10 was used in concentration 1.5 µg/µl, smaller particles resembling *T = 1* symmetry were identified. Similar smaller particles were also observed in samples after native virus purification [81]. The particle size can be influenced by the encapsulated NA, affecting the capsid structure and, subsequently, the size of nanoparticles [78, 126]. Additionally, the size of other components used for packaging can modulate the nanoparticle size depending on the molecular range [127].

**Figure 8.**
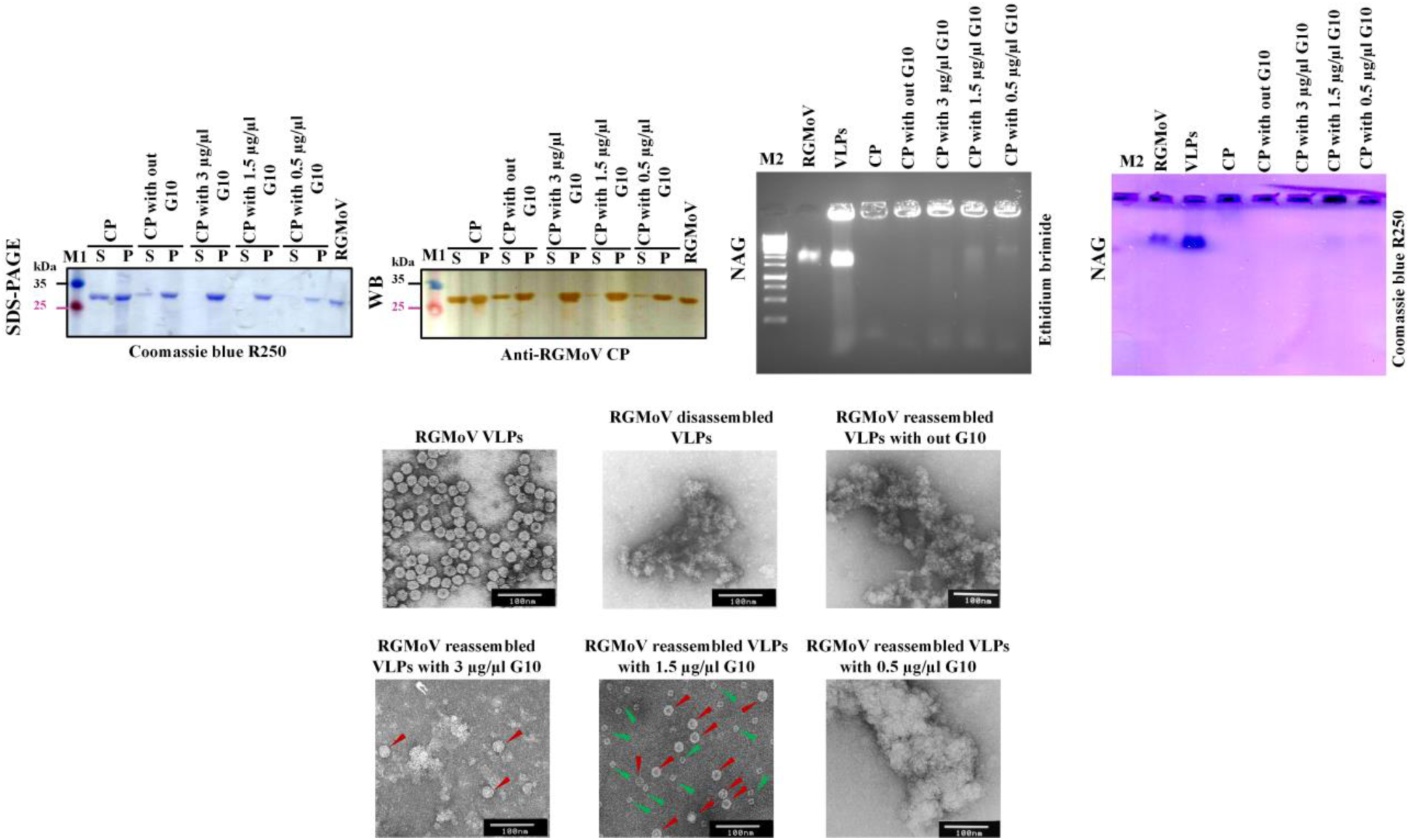
RGMoV CP derived VPL disassembly and reassembly analysis. SDS-PAGE – 12.5% dodecyl sulfate–polyacrylamide gel electrophoresis and gel staining with Coomassie blue R250 stain; WB – Western blot (primary antibodies: anti-RGMoV CP (1:1000); secondary antibodies: horseradish peroxidase-conjugated anti-rabbit IgG (1:1000, Sigma-Aldrich, USA); NAG – 0.8% native agarose gel visualized by ethidium bromide or Coomassie blue R250 TEM – transmission electron microscopy images with 1% uranyl acetate negative stain; red arrow indicate VLPs with *T = 3* symmetry; green arrow indicates VLPs with *T = 1* symmetry. S – soluble fraction after ultracentrifugation; P – solubilized pellet after ultracentrifugation; M1 – PageRuler™ Prestained Protein Ladder, 10-250 kDa (Thermo Fisher Scientific, USA); M2 – DNA 1 kb size marker (Thermo Fisher Scientific, USA).

The possible structural polymorphisms can be utilized to study the impact of different VLP sizes on the immune response regarding the time of target antibody identification, avidity, and neutralization level. Different forms of VLPs and their impact on the immune response have been studied, including VLPs with shape modifications through antigen insertion [22] or thermal remodeling [128, 129].

## 3. Conclusions

In conclusion, we demonstrated successful RGCP self-assembly into VLPs by transitioning from an episomal to a chromosome-integrated yeast expression system. The obtained VLPs resembled the native virus in both structure and stability. Additionally, the main encapsulated RNA originated from CP mRNA, demonstrating viral sequence structural compactness and selectivity. Using the BacterioMatch II Two-Hybrid system, we showed that RGCP has a higher binding capability to ssDNA compared to CCP and RCP. The RGCP’s ability to bind DNA can be utilized for applications such as CpG encapsulation, serving as a nanocontainer for NA delivery or as a vaccine platform for immune system modulation. These findings highlight the challenges of expressing and assembling viral CPs in different systems and underscore the need for optimized conditions tailored to each CP and expression host.

## 4. Materials and methods

### 4.1. Construction of expression systems

#### 4.1.1. Expression vector construction for *Escherichia coli* system

The construction of plasmids containing CfMV and RYMV *CPs* has been previously described [23]. For the RGMoV, we obtained the *CP* coding sequence from a plasmid containing the full-length cDNA (GenBank accession number: EF091714 [81, 130]) through polymerase chain reaction (PCR). For the RGMoV *CP*, we used primers RGCP-Nco-F and RGCP-Hind-R (Table 3), which contained NcoI and HindIII restriction sites. The PCR was performed on a Veriti 96 Well Thermal Cycler (Thermo Fisher Scientific, USA) using Pfu DNA polymerase (Thermo Fisher Scientific, USA) according to the manufacturer’s protocol, with a primer annealing temperature of 55 °C. The PCR product encoding the RGMoV *CP* was purified from agarose gel using the GeneJET Gel Extraction Kit (Thermo Fisher Scientific, USA) and subsequently cloned into the helper vector pTZ-57R/T using the InsTAclone PCR Cloning Kit (Thermo Fisher Scientific, USA), following adenine overlap addition by Taq polymerase (Thermo Fisher Scientific, USA) [72]. The ligation reaction was incubated at 65 °C for 10 min before transformation into XL1-Blue Supercompetent cells (Agilent Technologies, USA). Clones were selected via test restriction analysis on a 0.8% native agarose gel (NAG). At least three clones containing the *CP* insert were sequenced using M13-F and M13-R primers (Table 3), the ABI PRISM BigDye Terminator v3.1 Ready Reaction Cycle Sequencing Kit (Thermo Fisher Scientific, USA), and an ABI PRISM 3130xl sequencer (Thermo Fisher Scientific, USA). The *CP*-containing helper vector and the expression vector pET-28a(+) (Agilent Technologies, USA) were digested with NcoI (partial digestion for RGMoV *CPs*) and HindIII. After ligation, this resulted in the plasmid pET-RGCP.

For the expression vector containing the RGMoV *CP* with an N-terminally located six-histidine tag (N6H-RGCP), primers RGCP-Nde-F and RGCP-Hind-R (Table 3) were used. These primers contain the restriction sites NdeI and HindIII, respectively, and the pET-RGCP plasmid served as the template in PCR reaction. The cloning, selection and verification procedures for the *N6H-RGCP* insert-containing helper vector followed as previously described. The *N6H-RGCP* insert was then cloned into the pET-28a(+) vector between the NdeI and HindIII sites, resulting in the plasmid pET-N6H-RGCP.

#### 4.1.2. Vector construction for BacterioMatch II Two-Hybrid system

The *CPs* from CfMV, RYMV, and RGMoV were amplified and cloned into the pTRG vector from the BacterioMatch II Two-Hybrid system (Agilent Technologies, USA) using the following procedures. For CfMV *CP* (CCP), the amplification was carried out using primers Cf-CP-2Hyb-EcoRI-F and Cf-CP-2Hyb-BcuI-R resulting in a PCR product that contained BamHI and EcoRI restriction sites for subsequent cloning. Similarly, for RYMV *CP* (RCP), the amplification utilized primers – RY-CP-2Hyb-EcoRI-F and RY-CP-2Hyb-BcuI-R, producing a PCR product with EcoRI and BcuI restriction sites. For RGMoV *CP* (RGCP), the primers RG-CP-2Hyb-EcoRI-F and RG-CP-2Hyb-BcuI-R (Table 3) were used, generating a PCR product with EcoRI and BcuI restriction sites. The PCR products for each *CP* were then cloned into the pTZ-57R/T. The cloning strategy, selection and verification of positive clones followed previously described methods. Each *CP* were then cloned into the pTRG vector using the specified restriction sites. This process resulted in the successful creation of the pTRG-CCP vector for CfMV *CP*, the pTRG-RCP vector for RYMV *CP*, and the pTRG-RGCP vector for RGMoV *CP*.

#### 4.1.3. Expression vector construction for *Saccharomyces cerevisiae*

The RGMoV *CP* gene was amplified by PCR using primers RG-sc-F and RG-sc-R (Table 3). The PCR product of the *CP* gene was cloned, and the selection and verification strategy followed the previously described methods (see section 4.1.1). The RGMoV *CP* gene from the helper vector was cloned into the *S. cerevisiae* expression plasmid pFX7 between restriction sites XbaI and BglII. pFX7 carries the formaldehyde resistance gene for expression clone selection in yeast and the ampicillin resistance gene (*AmpR*) as a selective marker for plasmid selection in *E. coli* [74]. This cloning process resulted in the construction of the pFX-RGCP-sc plasmid.

#### 4.1.4. Plasmid construction for *P. pastoris* expression system

For the *P. pastoris* expression system, the RGMoV *CP* gene was amplified by PCR from pET-RGCP using primers RGCP-Pich-BamHI-F and RGCP-Pich-EcoRI-R (Table 3), which contain BamHI and EcoRI cloning sites. The *CP* gene from the helper vector was then cloned into the pPIC3.5K vector, which includes *AmpR* and gentamicin resistance (*GmR*) genes and is driven by the *AOX1* promoter (Thermo Fisher Scientific, USA). Following the cloning procedure, the resulting plasmid was named pPIC-RGCP. Prior to electroporation into *P. pastoris* competent cells, the pPIC-RGCP plasmid was digested with Ecl136II at the *AOX1* promoter region to linearize the plasmid. The linearized plasmid was then transfected by electroporation into *P. pastoris* cells along with ballast RNA to enhance transformation efficiency.

### 4.2. Expression and purification of CfMV, RYMV and RGMoV CPs

The coding sequence of the CfMV *CP* contains pool of rare tRNAs. To optimize expression, we used the BL21-CodonPlus (DE3)-RIL strain (Agilent Technologies, USA), which contains extra copies of the *argU*, *ileY*, and *leuW* tRNA genes. To avoid potential biases from the helper plasmid, both the control and all recombinant sobemovirus CPs were expressed in this strain. The expression plasmids pET-CCP, pET-RCP, pET-RGCP, and the empty pET-28a(+) were transformed into the BL21-CodonPlus (DE3)-RIL strain. For the pET-N6H-RGCP construct, the BL21(DE3) strain without a helper plasmid was used. Recombinant *E. coli* cells for CP expression were cultivated following an established protocol used for multiple plant virus CP expressions [23, 25, 36, 71]. After 16 h of expression at 20 °C, cells were harvested by low-speed centrifugation at 8000 rpm (8228 × g) for 5 min at 4 °C, weighed, and stored at -70 °C. The *E. coli* cells expressing corresponding CP were thawed on ice, resuspended in disruption buffer (15 mM KH_2_PO_4_ pH 5.5, 5 mM β-mercaptoethanol (β-ME), and disrupted by ultrasound using an ultrasound disintegrator UP200S (Hielscher Ultrasonics, Germany) at a period of 0.5 and intensity of 70% for 16 min. The protein soluble fraction was separated from the pellet by centrifugation at 11,000 rpm (15,557 × g) for 10 min at 4 °C.

*S. cerevisiae* strains AH22 *MATa leu2 his4* and DC5 *MATa leu2 his3* [131, 132] were transformed using the standard lithium acetate/PEG protocol [133]. Transformed yeast expression clones were selected on YPD agar (2% (w/v) peptone, 1% (w/v) yeast extract, 2% (w/v) glucose, 2% (w/v) agar) supplemented with 20 mM formaldehyde and cultivated as described by Sasnauskas et al. [134]. Expression was performed as previously described [76]. Cells were harvested by centrifugation at 8000 rpm (8228 × g) for 5 min at 4 °C, weighed, and stored at -70 °C. Yeast cells were thawed on ice, resuspended in disruption buffer (10 mM TRIS-HCl pH 7.0, 50 mM NaCl, 1 mM PMSF; 4 ml per 1 g of cells), and disrupted with a French press (Thermo Fisher Scientific, USA) in three cycles at 20,000 psi. The protein soluble fraction was separated from the pellet by centrifugation at 11,000 rpm (15,557 × g) for 10 min at 4 °C.

For the *P. pastoris* GS115 *his4* strain (Thermo Fisher Scientific, USA), plasmid electroporation, clone selection, and expression were conducted as previously described [75, 76]. Cells were harvested by centrifugation at 8000 rpm (8228 × g) for 5 min at 4 °C, weighed, and stored at -70 °C. Harvested *P. pastoris* biomass stored at -70 °C was thawed on ice and resuspended in disintegration buffer (15 mM KH_2_PO_4_ pH 5.5, 150 mM NaCl, 0.1% TX-100, 0.1% PMSF; 1 ml per 1 g of cells). Glass beads (acid-washed, 425-600 µm; Sigma-Aldrich, USA, Cat. No. G8772-100G) were added at 1/5 of the volume, and cells were disrupted using an ultrasound S220 device (Covaris, USA) at a peak intensity of 500, a duty factor of 20, and 500 cycles for 30 min at 7 °C. The protein soluble fraction was separated from the pellet by centrifugation at 11,000 rpm (15,557 × g) for 10 min at 4 °C.

Collected supernatants containing soluble CPs were purified on a sucrose dense gradient with five density steps (20%, 30%, 40%, 50%, 60% sucrose in buffer (15 mM KH_2_PO_4_ pH 5.5, 5 mM β-ME, 1% TX-100)). The gradient was performed in 36 ml open-top, thin-wall polyallomer tubes (Beckman Coulter, USA) in an ultracentrifuge Optima L-100 XP (Beckman Cultures, USA) with a swing-out rotor SW-32 at 25,000 rpm (106,559 × g) for 6 h at 18 °C. Preparation and sample removal after the sucrose gradient followed previously described methods [23].

After the sucrose gradient, fractions were analyzed by 12.5% SDS-PAGE and Western blotting (WB). Fractions containing CPs were pooled and dialyzed in a 12–14 kDa dialysis membrane (Spectrum Laboratories, USA) in a 1:100 dilution of dialysis buffer (15 mM KH_2_PO_4_ pH 5.5). After dialysis, CPs were concentrated in 26.3 ml polycarbonate ultracentrifuge tubes (Beckman Coulter, USA) in an ultracentrifuge Optima L-100 XP with a fixed angle rotor Type 70Ti (Beckman Coulter, USA) at 50,000 rpm (183,960 × g) for 4 h at 4 °C. Pellets were resuspended in 15 mM KH_2_PO_4_ pH 5.5, and an additional sucrose gradient was performed for higher VLP purity. Samples were measured on a NanoDrop-1000 (ND-1000; Thermo Fisher Scientific, USA) and a Qubit 2.0 (Thermo Fisher Scientific, USA) with a Qubit Protein Assay Kit (Thermo Fisher Scientific, USA). The codon optimized BL21-CodonPlus (DE3)-RIL strain and empty pET-28(a)+ vector (without sobemovirus *CPs*) served as a negative control in all CP expression and purification steps.

*E. coli* cells expressing N6H-RGCP were thawed on ice, resuspended in 1xLEW buffer (Affymetrix, USA), and disrupted by ultrasound with an ultrasound disintegrator UP200S (Hielscher Ultrasonics, Germany) at a period of 0.5 and intensity of 70% for 16 min. The protein soluble fraction was separated from the pellet by centrifugation at 11,000 rpm (15,557 × g) for 10 min at 4 °C. The pellet was dissolved in denaturation buffer according to the PrepEase™ Ni-IDA kit manual (Affymetrix, USA), and N6H-RGCP was purified on PrepEase™ Ni-IDA columns (Affymetrix, USA) under denaturing conditions. Elution fractions containing N6H-RGCP were refolded as described previously [135]. Protein purification steps were analyzed by 12.5% SDS-PAGE and WB with the His-Tag Antibody HRP Conjugate Kit (Merck-Millipore, Germany). Protein concentration was determined using the Bradford method (Bio-Rad, USA).

### 4.3. Plasmid analysis during CP expression

All purified CP samples (filtered through a 0.22 µm filter) from *E. coli* were transformed into XL1-Blue competent cells (Agilent Technologies, USA) to test for contaminant plasmids. The cells were grown on LB agar plates supplemented with kanamycin (Km, 25 µg/ml) and chloramphenicol (CAM, 25 µg/ml).

Samples for plasmid DNA extraction were collected at three time points, with 1 ml of cell culture per sample: from the start culture (T_S_), from non-induced culture directly before induction (OD600 0.8; T_0_), and from induced culture after 20 h of expression (T_Ex_). Plasmid DNA extraction was performed as previously described [136] with minor modifications. Plasmid DNA concentration was measured on a NanoDrop-1000 spectrophotometer at a wavelength of 260 nm. For *CP* identification in plasmids, equal amounts of plasmid DNA (20 ng/µl) were used for PCR reactions to test the plasmid for the *CP* gene. PCR was performed using a Veriti 96 Well Thermal Cycler. Taq polymerase was used according to the manufacturer’s protocol with primers pET-220 and pET-R (Table 1). PCR samples were analyzed on a 0.8% NAG.

### 4.4. Bacterial two hybrid system test

The XL1-Blue Kanamycin (Agilent Technologies, USA) strain was used for the BacterioMatch II Two-Hybrid system (Agilent Technologies, USA) to purify plasmids and detect CP interactions with nucleic acids (NA). The conditions for plasmid purification and interaction experiments were carried out as recommended by the manufacturer’s protocol. The *CP* genes of CfMV, RYMV, and RGMoV, cloned into the pTRG plasmid, were tested for interaction with NA using the BacterioMatch II Two-Hybrid system (Agilent Technologies, USA) with an empty pBT vector as a control. These experiments followed the procedures provided in the manufacturer’s protocol to ensure accurate detection of CP-NA interactions.

### 4.5. Sample analysis with SDS-PAGE, Western blotting (WB) and native agarose gel (NAG)

Proteins were fractionated on 12.5% SDS-PAGE according to the Laemmli [137] method and visualized using Coomassie brilliant blue R and/or G250 dyes (Sigma-Aldrich, USA). For WB analysis, proteins from the SDS-PAGE were transferred to a 0.45 µm nitrocellulose membrane (GE Healthcare, USA) at a current of 1 mA per cm² of SDS-PAGE for 45 min using the IMM-1-A semi-dry blotting system (The W.E.P. Company, USA). The CPs of RGMoV, CfMV, and RYMV were detected on the membrane using specific rabbit polyclonal antibodies. These antibodies included anti-RGMoV (produced in-house from the native virus purified from *Avena sativa* [34, 81]), anti-CfMV (produced in-house from the native virus purified from *A. sativa* [33]), and anti-RYMV (produced in-house from VLPs purified from *E. coli* [23]). The antibodies were used at a 1:1000 dilution. The secondary antibody was an anti-rabbit IgG (whole molecule) peroxidase conjugate (Sigma-Aldrich, USA), also diluted 1:1000. Visualization of the bands was carried out using the substrate o-dianisidine (0.002% (w/v), Sigma-Aldrich, USA) with 0.03% (v/v) H₂O₂, and the bands were detected by visual color development [81].

Samples containing CPs were mixed with 6x DNA loading dye (Thermo Fisher Scientific, USA) and loaded onto a 0.8% native agarose gel (NAG). Nucleic acids (NA) were detected by staining the NAG with ethidium bromide (0.5 mg/ml; Thermo Fisher Scientific, USA) and visualized under UV light. For protein visualization, the NAG was stained with Coomassie Brilliant Blue R and/or G250 dyes [23].

### 4.6. Analysis of CP containing samples by TEM, MS, DLS, Sypro-Orange dye, RT-PCR, RAP-PCR, and HTS

For TEM analysis, purified CPs and control samples were adsorbed onto carbon formvar-coated copper grids and negatively stained with a 1% uranyl acetate aqueous solutions as described previously [72]. The grids were examined using a JEM-1230 transmission electron microscope (TEM; JEOL, Japan) at an accelerating voltage of 100 kV.

MALDI TOF mass spectrometry analysis of RGMoV CP samples (1 mg/ml) isolated from oat plants [81] and *P. pastoris* was conducted according to the protocol described by Balke, Resevica et al. [23].

Dynamic Light Scattering (DLS) analysis for RGMoV VLPs and the native virus (1 mg/ml) was carried out as described in Balke, Resevica et al. [23] using a Zetasizer Nano ZS DLS instrument (Malvern Instruments Ltd., UK) and DTS software, version 7.11 (Malvern Instruments Ltd., UK).

To compare the stability of purified native RGMoV and its CP-derived VLPs under different temperatures and additives, structural changes were monitored in real-time using an MJ Mini PCR system (Bio-Rad, USA) with a DNA melting point determination program and Sypro-Orange dye (Sigma, USA), following methods described previously [25, 36, 100]. Data were analyzed using Opticon Monitor Software (Bio-Rad, USA).

All purified CP and control samples (1 mg/ml, regardless of the expression system) were treated with benzonase (25 U/µl; Novagen, Germany) to remove unprotected NA before RNA isolation. Encapsulated RNA was isolated using TRI REAGENT (Sigma-Aldrich, USA) per the manufacturer’s protocol and solubilized in 30 µl RT-PCR grade H_2_O (Thermo Fisher Scientific, USA). Concentration was determined using a NanoDrop-1000 and Qubit 2.0 with a Qubit RNA high sensitivity assay kit (Thermo Fisher Scientific, USA), and stored at -80 °C.

An adapted RNA arbitrarily primed polymerase chain reaction (RAP-PCR) [138] was performed to identify the genetic background of purified VLPs from *S. cerevisiae*. Firstly, an RT reaction was performed for RNA (0.5 µg) isolated from *S. cerevisiae* purified 40 nm VLPs using randomly selected primer from laboratory stock 99.R and RevertAid H Minus Reverse Transcriptase (Thermo Fisher Scientific, USA) performing cDNA synthesis step at 37 °C, all procedures were according to provided standard protocol to produce cDNA. Secondly a PCR was performed using the same primer 99.R (Table 3) and Taq polymerase (Thermo Fisher Scientific, USA) according to the manufacturer’s protocol, with a primer annealing temperature of 55 °C. RAP-PCR products were directly cloned into the previously used helper vector pTZ-57. Plasmids containing inserts, verified by restriction enzyme tests with EcoRI and HindIII, were sequenced with the M13-F primer by Sanger sequencing method (Table 3).

RT-PCR for detection of *CP*, *23S rRNA* and *16S rRNA* coding sequences from RNA isolated form *E. coli* purified samples for cDNA synthesis was performed with random hexamer primers (Thermo Fisher Scientific, USA) and RevertAid H Minus Reverse Transcriptase according to the provided protocol. For PCR reactions, Pfu polymerase (Thermo Fisher Scientific, USA) was used with the following primer sets: for CfMV *CP* (CCP-F and CCP-R, Table 3), RYMV *CP* (RCP-stop-R and RY-Ct1, Table 3), RGMoV *CP* (RG-T1-sc-F and RG-CP-seq-R, Table 3), plasmid-specific primers (pET-F and pET-R, Table 3), and *16S rRNA* (16sRNA-F and 16sRNA-R, Table 3) and *23S rRNA* (23sRNA-F and 23sRNA-R, Table 3). Direct PCR on purified *CPs* was performed with primers pET-F and pET-R (Table 3) and Pfu polymerase following the manufacturer’s protocol. All RT-PCR or PCR reactions were analyzed on 0.8% NAG.

High-throughput sequencing (HTS) libraries for RGMoV CP derived VLPs were prepared according to the Ion Total RNA-Seq Kit (Thermo Fisher Scientific, USA) protocol. HTS was performed on the Ion Torrent Personal Genome Machine™ (PGM™) sequencer using the Ion PGM Sequencing 200 Kit (Thermo Fisher Scientific, USA) according to standard protocols.

Detailed procedures are available elsewhere [81]. HTS libraries for CfMV and RYMV CP derived VLPs were prepared according to the Ion Total RNA-Seq Kit v2 (Thermo Fisher Scientific, USA). HTS was performed on the Ion Torrent Proton (Thermo Fisher Scientific, USA) using the Ion PI™ Hi-Q™ Sequencing 200 Kit (Thermo Fisher Scientific, USA) according to standard protocols.

### 4.7. HTS data analysis

For RNA-Seq library sequencing, reads were trimmed for adapters, barcodes, and quality using Cutadapt version 4.2 [139] and Fastp version 0.23.2 [140]. Reads with a minimum length of 50 bp were retained for further analysis using featureCounts version 2.0.0 [141] from the Subread bioinformatics tool package. The following parameters were used: for host origin reads: -’t gene -g Namè, for plasmid origin reads: the ‘-t’ parameter was used, with the specific feature types set accordingly. Plasmid .gb files were converted to gff3 format using the Galaxy "Genbank to GFF3" tool with default parameters. For the host genome, gff3 files were downloaded from: NC_012963.1, AM946981.2.

### 4.8. Genomic DNA isolation from *P. pastoris* for RGMoV CP gene detection

Genomic DNA from *P. pastoris* was isolated using the lithium acetate and NaOH method as previously described [142]. PCR was then performed with Taq polymerase and primers RGCP-Pic-BamHI-F and RGCP-Pich-EcoRI-R (Table 3) according to the manufacturer’s protocol. The PCR products were analyzed on a 0.8% NAG stained with ethidium bromide (0.5 mg/ml).

### 4.9. Gel-shift analysis for RGMoV CP binding ability to ss and dsDNA

Gel-shift assays for RGCP interaction with ss and dsDNA were carried out using a previously described method [136]. N6H-RGCP was analyzed in a 0.6% NAG with increasing concentrations of the CP while maintaining constant concentrations of ssDNA or dsDNA. The DNA source used was the pET-N6H-RGCP plasmid and the RGMoV *CP* gene PCR product. For assessing dsDNA, the pET-N6H-RGCP plasmid was linearized using HindIII or digested with KpnI and BglII. To obtain ssDNA, all dsDNA samples were thermally desaturated at 95 °C for 10 min. The ds or ssDNA with fixed concentrations (linearized pET-N6H-RGCP at 250 ng; double-digested pET-N6H-RGCP at 3.4 µg; RGMoV *CP* PCR product at 255 ng) was mixed with increasing concentrations of N6H-RGCP (175 µg; 250 µg; 500 µg; 1000 µg; 2000 µg; 4000 µg). The samples were incubated on ice for 10 min, then mixed with 5 µl of 6x DNA loading dye (Thermo Fisher Scientific, USA) and loaded onto a 0.6% NAG.

### 4.10. RGMoV CP derived VLP disassembly and *in vitro* reassembly

0.5 mg of purified RGMoV VLPs were disassembled ON at 4 °C in disassembly buffer (100 mM TRIS-HCl pH 8.0, 600 mM NaCl, 30 mM EDTA, 30 mM DTT, 1 mM PMSF) with a final reaction volume of 500 µl. Following disassembly, the CP was centrifuged at 72,000 rpm (280,000 × g) for 1 h at 5 °C in a TLA-100.3 rotor (Beckman Coulter, USA) on a TL-100 ultracentrifuge (Beckman Coulter, USA) to separate the disassembled CP molecules from oligomers or intact particles. The pellet was dissolved in 500 µl of 15 mM KH_2_PO_4_ pH 5.5, matching the volume used for disassembly, for analysis in 12.5% SDS-PAGE. Only the supernatant was used for reassembly experiments. The concentration of CP in the soluble fraction was measured using a ND-1000. For each reassembly test sample, 100 µl (120 µg) of disassembled CP sample was used. Four conditions were tested for VLP reassembly: 1) plain CP, 2) CP with 3 µg/µl of a type A CpG TLR9 agonist—G10 (5’ GGGGGGGGGGGACGATCGTCGGGGGGGGGG 3’), 3) CP with 1.5 µg/µl G10, and 4) CP with 0.75 µg/µl G10. *In vitro* assembly was performed by dialyzing the samples in 200 volumes of 20 mM TRIS-HCl pH 7.0, 1 mM PMSF, 2 mM CaCl_2_ buffer ON at 4 °C. G10 was added directly in CP sample. After in vitro reassembly, the samples were centrifuged at 72,000 rpm (280,000 × g) for 1 h at 5 °C in a TL-100.1 rotor (Beckman Coulter, USA) on a TL-100 ultracentrifuge to pellet the reassembled VLPs. The pellet was resuspended in 100 µl of 15 mM KHPO_4_ pH 5.5 and, after solubilization, was measured using the ND-1000 along with supernatant after ultracentrifugation. The pellet after solubilization and the supernatant after ultracentrifugation were analyzed by 12.5% SDS-PAGE and WB with anti-RGMoV antibodies. The pellet after solubilization also was analyzed by NAG and TEM.

## CRediT authorship contribution statement

**Ina Balke:** Conceptualization, Methodology, Data curation, Formal analysis, Visualization, Validation, Investigation, Supervision, Project administration, Funding acquisition, Writing – original draft, Writing – review & editing. **Gunta Resevica:** Investigation, Methodology, Data curation, Validation. **Vilija Zeltina:** Investigation, Methodology, Data curation, Validation. **Ivars Silamikelis:** Conceptualization, Methodology, Data curation, Formal analysis, Visualization, Validation, Investigation, Supervision, Writing – review & editing. **Elva Liepa:** Investigation, Methodology, Data curation, Validation. **Reinis Liepa:** Investigation, Methodology, Data curation, Validation. **Ieva Kalnciema:** Investigation, Methodology, Data curation, Validation. **Ilze Radovica-Spalvina:** Methodology, Data curation, Formal analysis, Validation, Investigation. **Dita Gudra:** Methodology, Data curation, Formal analysis, Validation, Investigation. **Janis Pjalkovskis:** Methodology, Data curation, Formal analysis. **Janis Freivalds:** Investigation, Methodology, Data curation, Validation. **Andris Kazaks:** Conceptualization, Methodology, Data curation, Formal analysis, Validation, Investigation, Supervision, Writing – review & editing. **Andris Zeltins:** Conceptualization, Methodology, Data curation, Formal analysis, Visualization, Validation, Investigation, Supervision, Writing – review & editing.

## Competing interests

The authors declare that they have no known competing financial interests or personal relationships that could have appeared to influence the work reported in this paper. The funders had no role in the design of the study; in the collection, analyses, or interpretation of data; in the writing of the manuscript; or in the decision to publish the results.

## Acknowledgements

We acknowledge the MAFF Genbank for the RGMoV isolate. We thank Dace Skrastiņa from the Latvian Biomedical Research and Study Centre’s Biomedical Technology Complex, Laboratory Animal Core Facility, for providing polyclonal antibodies against RGMoV, RYMV, and CfMV CPs. We also extend our gratitude to Davids Fridmanis from the Latvian Biomedical Research and Study Centre’s Bioinformatics Core Facility for his continuous support and advice in HTS data analysis methodology. Additionally, we appreciate the assistance of Janis Bogans from the Latvian Biomedical Research and Study Centre’s Biotechnology Core Facility in sample DLS analysis. We would like to thank Velta Ose for her invaluable contribution to conducting the TEM analysis of the purified virus and VLPs samples.

This research was funded by the Latvian Council of Science, Grant No. lzp-2022/1-0629.

## Data availability

Data will be made available on request.

## Declaration of generative AI and AI-assisted technologies in the writing process

During the preparation of this work the author(s) used ChatGPT in order to improve language and readability. After using this tool/service, the author(s) reviewed and edited the content as needed and take(s) full responsibility for the content of the publication.

## Notes

### Competing Interest Statement

The authors have declared no competing interest.

