## Supporting Information for "Investigation of plant virus-like particle formation in bacterial and yeast expression systems"

^e^“Exotic” site microbiome and G-protein coupled receptor functional research group, Latvian Biomedical Research and Study Centre, Ratsupites street 1, k-1, LV-1067, Riga, Latvia;

^f^Structural biology group, Latvian Biomedical Research and Study Centre, Ratsupites street 1, k-1, LV-1067, Riga, Latvia

**Supplementary Figures**

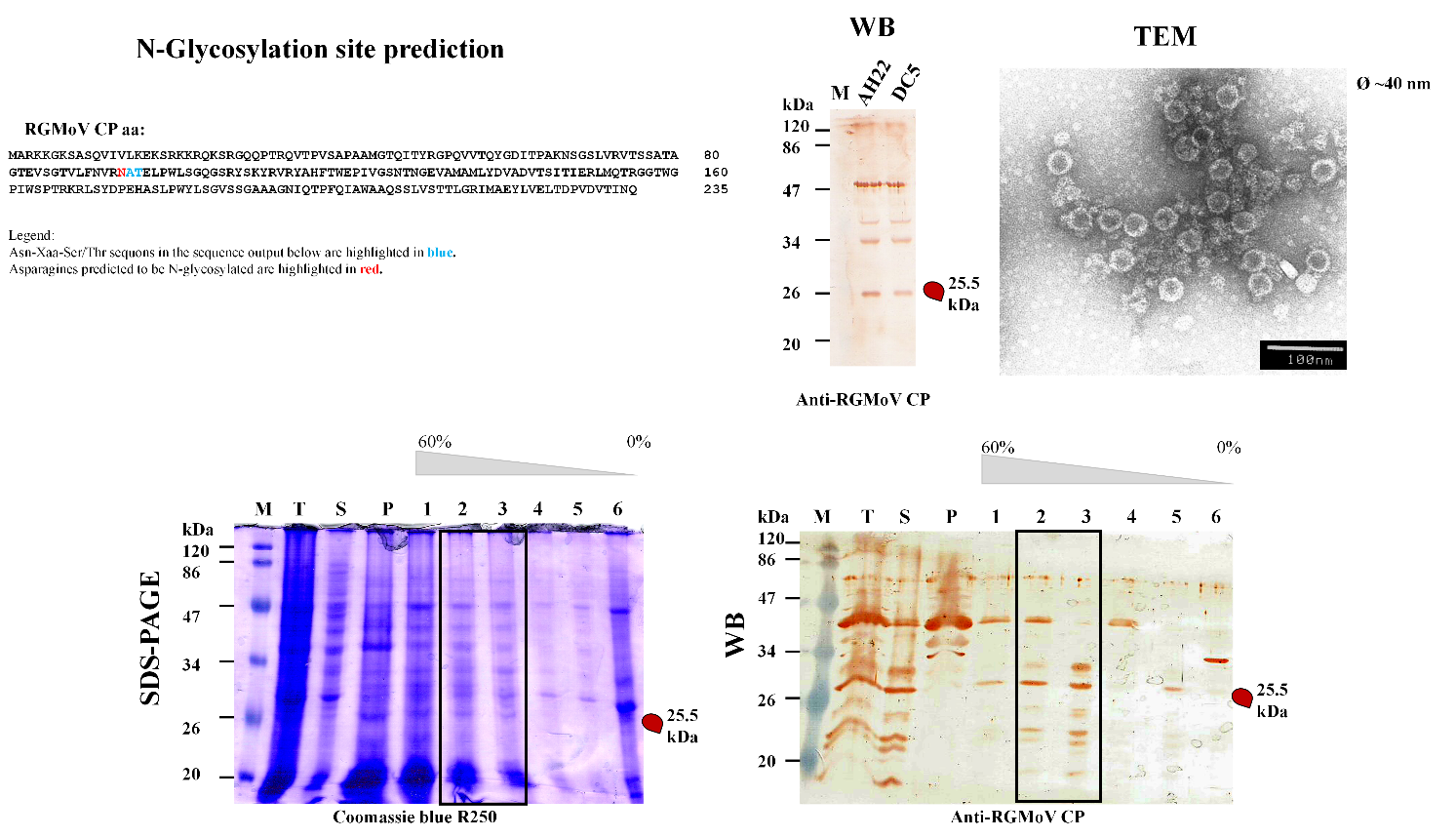

**Figure S1.** RGMoV CP amino acid sequence for N-glycosylation and expression in S. cerevisiae and purified CP analysis. M – pre-stained protein marker (Weight Marker, Thermo Fisher Scientific, USA); T – total cell lysate after expression; S – supernatant after cell French press and sample clarification; P – pellet after cell French press and sample clarification; 1 – sucrose gradient fraction containing 60% sucrose; 2 – sucrose gradient fraction containing 50% sucrose; 3 – sucrose gradient fraction containing 40% sucrose; 4 – sucrose gradient fraction containing 30% sucrose; 5 – sucrose gradient fraction containing 20% sucrose; 6 – top fraction after sucrose gradient; SDS-PAGE – sample analysis by 12.5% dodecyl sulfate–polyacrylamide gel electrophoresis and gel staining with Coomassie blue R250 stain; WB – Western blot (primary antibodies: anti-RGMoV CP (1:1000); secondary antibodies: horseradish peroxidase-conjugated anti-rabbit IgG (1:1000, Sigma-Aldrich, USA); TEM – transmission electron microscopy images with 1% uranyl acetate negative stain. SDS-PAGE and WB tracks marked in the rectangle were used for further purification. N-glycosylation analysis was performed with NetNGlyc - 1.0 [1].

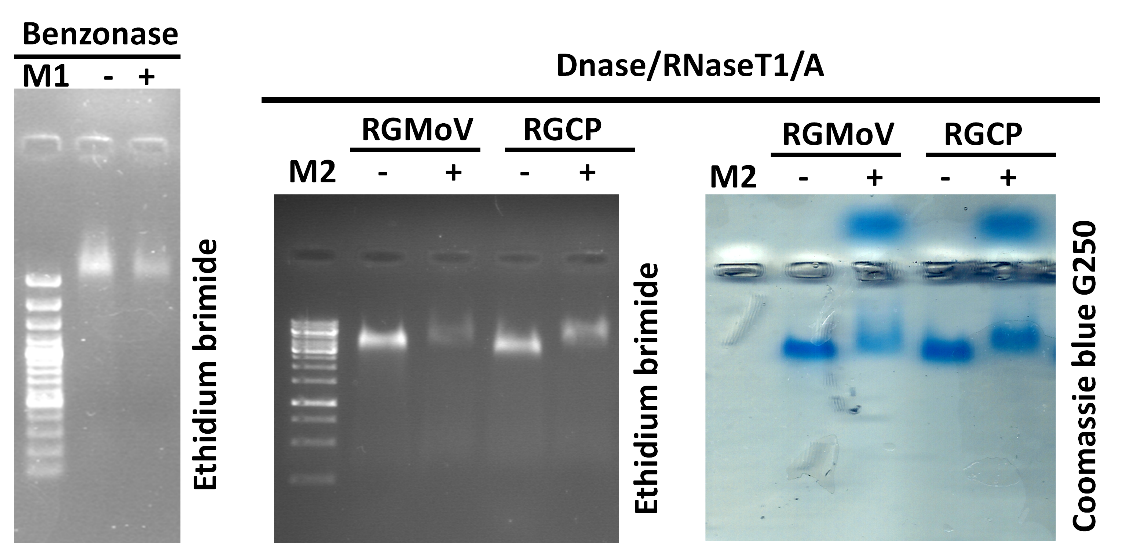

**Figure S2.** Nuclease resistance analysis. RGMoV – native virus purified from *A. sative* (MW – 25.6 kDa); 5 µg; RGCP – RGMoV CP derived VLPs purified form *P. pastoris* (MW – 25.6 kDa); 5 µg; M1 – GeneRuler 100 bp Plus DNA Ladder (Thermo Fisher Scientific); M2 - DNA 1 kb size marker (Thermo Fisher Scientific, USA); NAG – 0.8% native agarose gel; SDS-PAGE – 12.5% dodecyl sulfate–polyacrylamide gel electrophoresis.

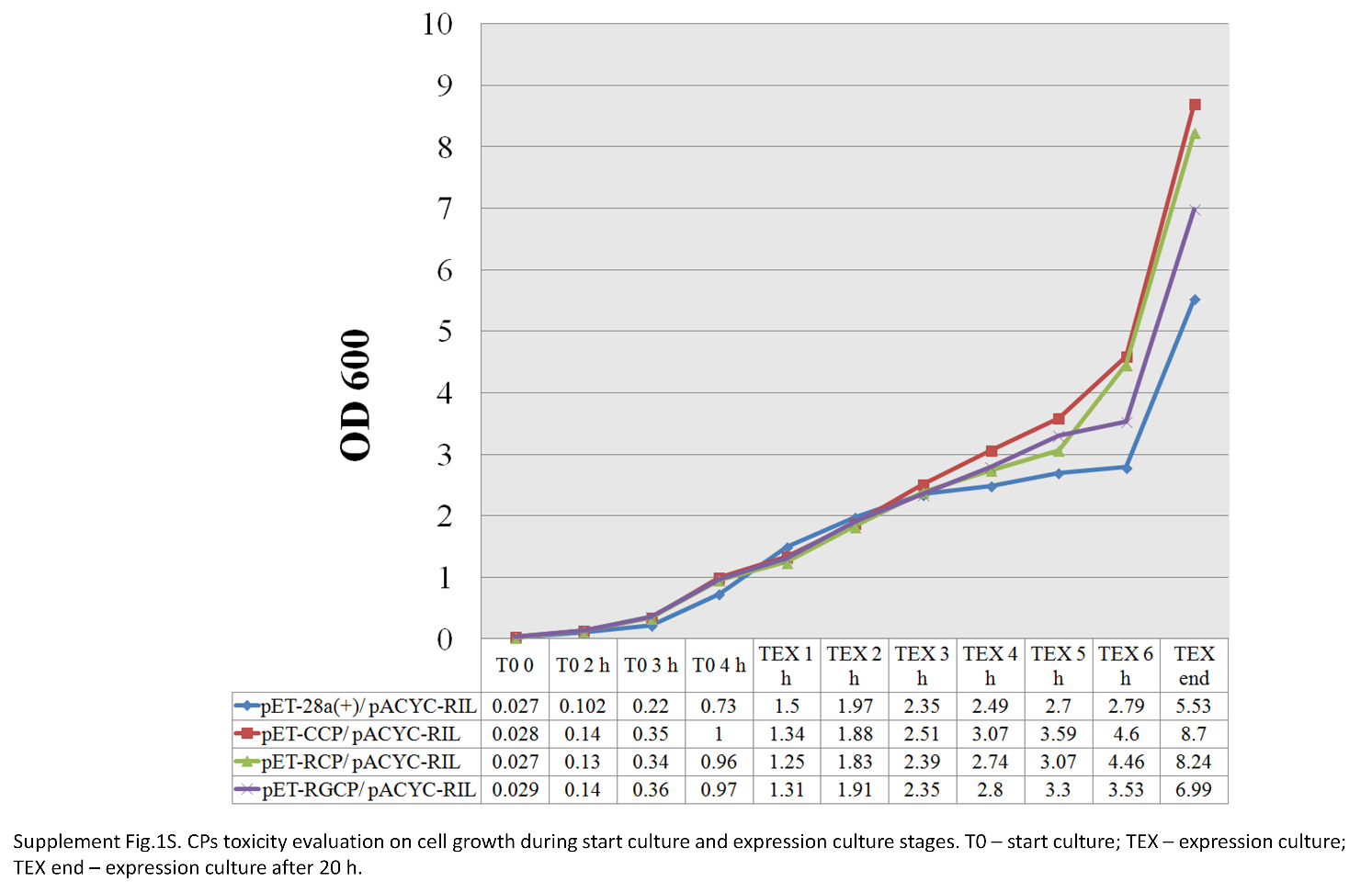

**Figure S3.** CPs toxicity evaluation on cell growth during start culture and expression culture stages. T_0_ – seed culture; T_EX_ – expression cultures at different time points after induction with 0.2 mM IPTG at OD_600_ 1 (1 h, 2 h, 3 h, 4 h, 5 h, 6h); T_EX end_ – expression culture after 20 h. Results represent the average value from three replicates.

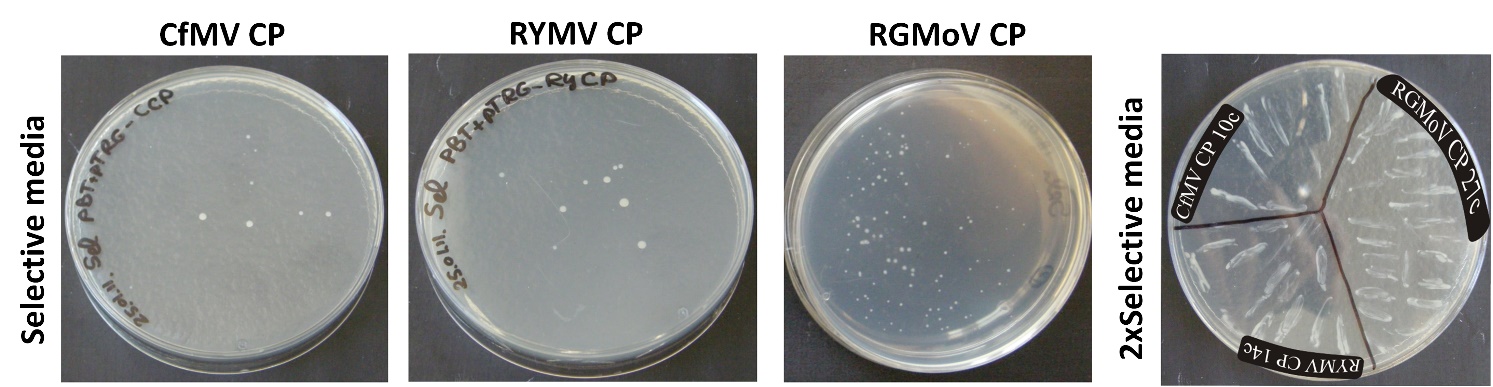

**Figure S4.** CfMV CP, RYMV CP and RGMoV CP nucleic acid binding test with BacterioMatch II Two-Hybrid system. 10c, 14c, 27c – colony number plated from plate with selective media to 2xSelective media.
